# Chronic α-Synuclein Over-Expression and Ceruloplasmin Challenge Promote Distinct Iron and Redox Responses in M17 Cells

**DOI:** 10.64898/2026.06.15.732494

**Authors:** Masoumeh Pourhadi, Bedri Ranxhi, Seth P. Sukuria, Fatimah H. Hussein, Sokol V. Todi, Peter A. LeWitt, Wei-Ling Tsou

## Abstract

**Background:** While α-synuclein (α-syn) accumulation and iron dysregulation are hallmarks of Parkinson’s Disease, the adaptations that enable neuronal survival under chronic protein stress remain unclear. Here, we investigated how α-syn overexpression and ceruloplasmin (Cp)-mediated iron modulation alters iron and redox homeostasis.

**Methods:** We utilized human BE(2)-M17 neuroblastoma cell lines stably expressing different levels of α-syn to examine the interplay between α-syn, Ceruloplasmin (Cp)-mediated iron modulation, and the cellular response to oxidative stress. Analyses included Western blotting, immunofluorescence staining, soluble/insoluble fractionation, glutathione, reactive oxygen species (ROS) and hydrogen peroxide (H₂O₂) quantification, lipid peroxidation, ferrous iron, and cell viability.

**Results:** Our data suggest an unexpected relationship between chronic α-syn expression and cellular redox regulation. Despite carrying a greater α-syn burden, cells with higher α-syn expression exhibit lower basal ROS, H₂O₂, and lipid peroxidation compared to control cells. These changes are not accompanied by activation of canonical antioxidant pathways suggesting that the reduced oxidative profile arises through alternative mechanisms. Besides, α-syn over-expressing cells display significant remodeling of iron-handling pathways, including altered expression of ferritin heavy chain, transferrin receptor, and ferroportin, suggesting that chronically high α-syn levels are associated with changes in iron homeostasis. In addition, this phenotype is not maintained following Cp overexpression. Although Cp reduces Fe²⁺ levels, it also induces substantial increases in ROS and H₂O₂ without corresponding changes in GPX4, glutathione, or related antioxidant systems. Thus, the reduced basal oxidative profile observed in α-syn-over-expressing cells does not reflect enhanced canonical antioxidant capacity. Instead, chronically high α-syn levels appear to be associated with adaptive remodeling of iron and redox pathways that become sensitive to oxidative imbalance.

**Conclusion:** Chronic α-syn over-expression promotes adaptive remodeling of iron and redox homeostasis, associated with reduced basal oxidative stress but increased sensitivity to Cp-mediated perturbation. These data link α-syn burden to iron metabolism and stress-dependent vulnerability in synucleinopathies.

**Graphical abstract:** 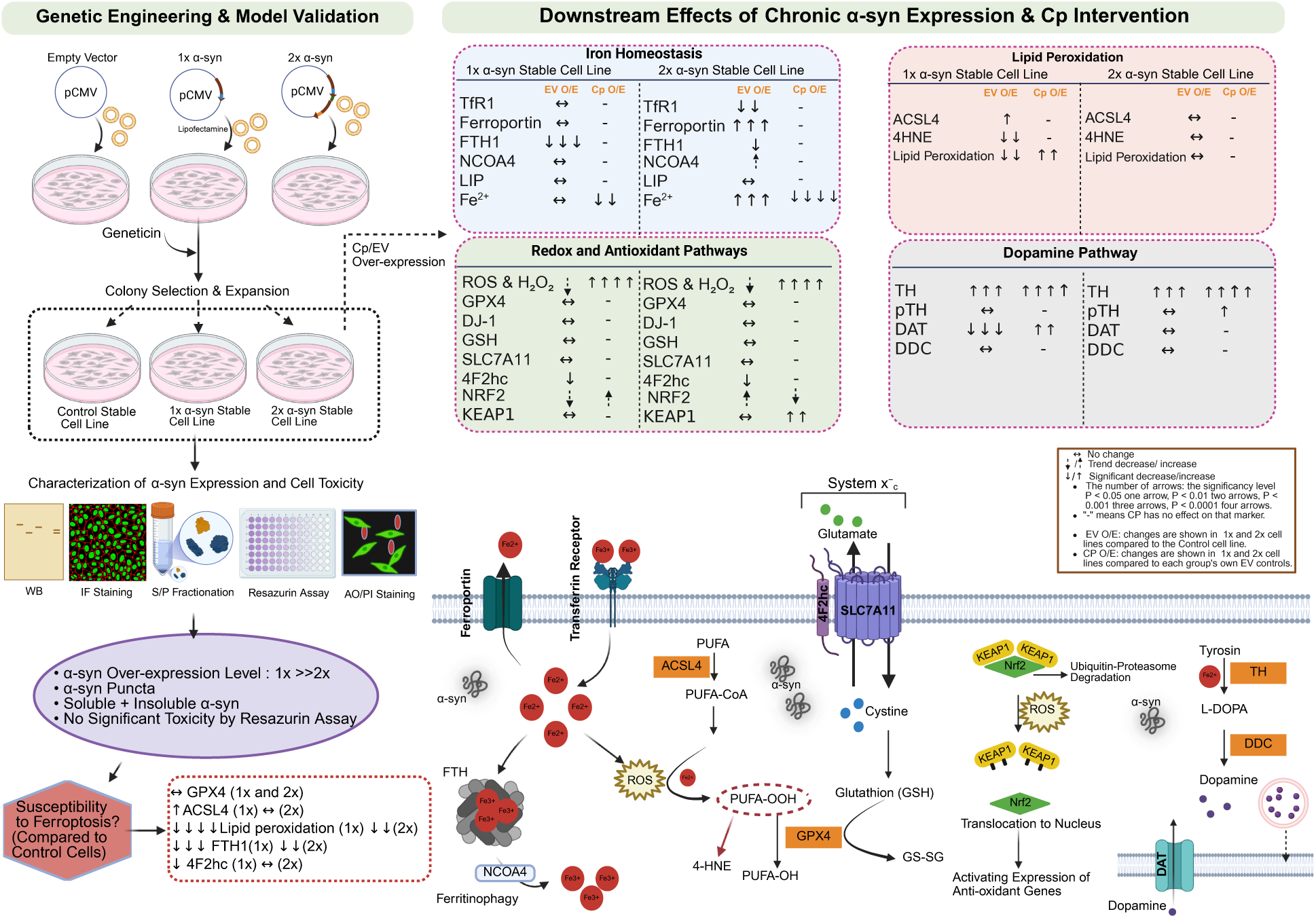

## 1 INTRODUCTION

Parkinson’s disease (PD) is a progressive neurodegenerative disorder whose motor features are largely attributable to the continuing loss of dopaminergic neurons in the substantia nigra pars compacta (SNpc) (1). A central contributor to this process is α-synuclein (α-syn), a synaptic protein present in the SNpc that can misfold, accumulate, and transition from soluble species into amyloid-rich aggregates that are characteristic of PD neuronal pathology. Although α-syn accumulation is a defining end-stage molecular feature of the disease, the chronic mechanisms by which chronic α-synuclein burden builds up and disrupts cellular homeostasis remain incompletely understood (2, 3).

One major cellular pathway linking α-syn toxicity to neuronal dysfunction is the disruption of iron and redox balance. Cationic iron is an essential factor in neuronal metabolism. It is particularly important for dopaminergic neurons because ferrous iron (Fe²⁺) is a component of tyrosine hydroxylase (TH), the rate-limiting enzyme in dopamine biosynthesis. However, excess labile Fe²⁺ in cells can also be toxic. Through the Fenton reaction, Fe²⁺ promotes the generation of reactive oxygen species (ROS), which can overwhelm antioxidant defenses like glutathione and drive formation of lipid peroxides. This form of membrane oxidative damage is also closely associated with ferroptosis, an iron-dependent cell death pathway increasingly implicated in the neuronal loss of PD and for related neurodegenerative disorders (4, 5).

Ceruloplasmin (Cp) is an important regulator of the iron-redox interface. As a ferroxidase, Cp converts reactive Fe²⁺ into ferric iron (Fe³⁺), facilitating export of Fe^3+^ through ferroportin and limiting the accumulation of redox-active iron (6). For this reason, Cp can be considered neuroprotective (7, 8). However, its role may be more complicated in the setting of a chronic α-syn burden, lie PD and other synucleinopathies. Although reduced Cp activity has been associated with redox-active iron accumulation in brain, it is not known if increasing Cp levels restores iron and redox balance in α-syn-stressed cells, particularly if these cells have already diminished the antioxidant plasticity hampering their ability for adapting to additional stress (7, 9).

This uncertainty reflects a broader paradox in α-syn biology. While various models of neurodegeneration predict that increasing α-syn burden directly enhances oxidative stress and cellular dysfunction, emerging evidence also suggests that chronic protein accumulation may also activate adaptive proteostasis and stress-response pathways that help maintain cellular viability despite persistent proteotoxic challenges. (10, 11). Such adaptations may involve alterations in iron handling, redox regulation, and lipid metabolism, potentially reshaping how cells respond to subsequent stressors. As a result, cellular vulnerability may not be determined solely by the magnitude of α-syn protein levels, but also by the nature of the adaptive responses that develop over time. Understanding these responses may provide insight into why some cells remain functionally stable despite chronic α-syn burden, whereas others become increasingly susceptible to secondary insults (12–14).

In this study, we examined how stable α-syn over-expression alters cellular responses to iron and oxidative stress in immortalized BE(2)-M17 neuroblastoma cells. Specifically, we investigated (1) how α-syn over-expression influences iron import, storage, export, and the labile iron pool; (2) how antioxidant signaling pathways, including the Nrf2/KEAP1 axis and System Xc^-^ respond to Cp-mediated perturbation; and (3) how these changes affect lipid peroxidation and dopaminergic markers, including TH, dopa decarboxylase (DDC), and dopamine transporter (DAT). Our studies were designed to explore how chronically high α-syn protein levels remodel iron homeostasis, redox regulation, and dopaminergic pathways, and whether these adaptations influence cellular responses to secondary intervention. We found that chronic α-syn over-expression is associated with altered iron handling, reduced basal oxidative stress, and changes in lipid peroxidation and dopaminergic signaling. Furthermore, Cp exposure reveals distinct responses among the α-syn-expressing cell lines, suggesting that chronic α-syn burden promotes adaptive remodeling of cellular homeostasis while simultaneously creating context-dependent vulnerabilities that become apparent under secondary stress.

## 2 Materials and Methods

### 2-1 Generation of Stable α-Synuclein–Expressing BE(2)-M17 Cell Lines

Human BE(2)-M17 neuroblastoma cells (ATCC; from here on referred to as M17 cells) were cultured in a 1:1 mixture of Eagle’s Minimum Essential Medium and Ham’s F-12 (EMEM/F12; ATCC), supplemented with 10% fetal bovine serum (FBS; Gibco) and 1% penicillin–streptomycin (Gibco). Cells were maintained at 37°C in a humidified incubator with 5% CO₂ and passaged at 70–80% confluency using 0.25% trypsin–EDTA. Cells were routinely tested for mycoplasma contamination and confirmed to be negative.

Human α-syn (SNCA) cDNA was synthesized and cloned into pCMV-3Tag-3A-P2A mammalian expression vector by GenScript to generate one copy α-syn plasmid (1x-α-syn). A two-copy α-syn construct (2x-α-syn) was generated by inserting a second α-syn coding sequence downstream of a self-cleaving P2A peptide, enabling co-expression of two proteins from a single transcript. The upstream α-syn has a C-terminal HA epitope tag, while the downstream α-syn has a C-terminal FLAG tag.

For generation of stable lines, cells were transfected with plasmids encoding either 1x-α-syn or 2x-α-syn, or empty vectors (EV) as a Control. Cells were seeded at 2.5 × 10⁵ cells per well in 12-well plates and maintained in antibiotic-free medium prior to transfection. Transfection was performed using Lipofectamine LTX with PLUS reagent (Thermo Fisher Scientific). Briefly, 1 µg plasmid DNA was mixed with 1 µL PLUS reagent in Opti-MEM (Thermo Fisher Scientific), followed by addition of 3 µL Lipofectamine LTX. Complexes were incubated for 20 minutes at room temperature and then added to cells. After 24 hours, medium was replaced, and selection was initiated using Geneticin (G418; 900 µg/mL, Gibco). After approximately two weeks, resistant colonies were expanded and maintained in 400 µg/mL G418. Stable expression was confirmed by Western blotting and immunofluorescence staining.

### 2-2 Cell Viability Assays

#### 2-2-1 Resazurin Reduction Assay

Cell metabolic activity was assessed using a resazurin-based assay (Resazurin Cell Viability Kit, Cell Signaling). Control, 1x, or 2x α-syn stable cell lines were seeded in 96-well plates at densities of 2.5×10³ or 8×10³ cells per well and allowed to adhere overnight and Medium was replaced as needed during the experimental period. Resazurin solution (10% v/v of well volume) was added, and cells were incubated for 3 hours at 37°C. Metabolically active cells reduce resazurin to resorufin, resulting in a fluorescent signal. Fluorescence was measured using a Tecan Infinite M Plex plate reader (excitation: 560 nm; emission: 590 nm). Cell viability was expressed as a percentage relative to Control cells on the same day.

#### 2-2-2 AOPI Staining

Cell viability was assessed using acridine orange/propidium iodide (AOPI) staining. Control, 1x, and 2x α-syn cell lines were seeded at 3 × 10⁴ cells per well in 12-well plates and stained with ViaStain AO/PI solution (2 µg/mL, Revvity) for 2–5 minutes at room temperature, protected from light. Cells were imaged using EVOS M5000 microscope (Invitrogen). Live cells were identified by acridine orange staining (green fluorescence), whereas dead cells were identified by propidium iodide staining (red fluorescence). The percentage of viable cells was calculated as the ratio of live cells to total cells using Image J software. During the 8-day experimental period, the culture medium was not removed; instead, fresh medium was added as needed.

### 2-3 Soluble and Insoluble Fractionation

To separate proteins into soluble and insoluble fractions, Control, 1x, and 2x α-syn cells were washed with ice-cold phosphate-buffered saline (PBS; Gibco) and lysed in NETN buffer (50 mM Tris-HCl, pH 7.5, 150 mM NaCl, 0.5% Nonidet P-40) supplemented with 1 mM PMSF, protease inhibitor cocktail III (Calbiochem), and Halt phosphatase inhibitor cocktail (Thermo Scientific). Lysates were briefly sonicated on ice (50% amplitude, 15 s) and centrifuged at 20,000 × g for 30 minutes at 4°C. The supernatant was collected as the soluble fraction. The remaining pellet was washed once with NETN buffer, briefly vortexed, and centrifuged again under the same conditions. After removing the supernatant, the pellet (insoluble fraction) was resuspended in 50 µL Laemmli buffer (12.5% 1 M Tris, 20% glycerol, 4% SDS, pH 7.5). To ensure complete solubilization, the samples were further sonicated (50% amplitude, 30 s). Both soluble and insoluble fractions were then boiled at 95°C for 5 minutes and used for subsequent Western blot analysis.

### 2-4 Transient Over-expression of Ceruloplasmin

Human Cp cDNA encoding the full-length human Cp protein (∼1046 amino acids) was cloned into a pCMV-based mammalian expression vector with 3-MYC and FLAG tags to enable constitutive expression by GeneScript. The Control, 1x, and 2x α-syn cell lines were transfected with either Cp plasmid (pCMV-Cp) or empty vectors (EV, pCMV) using Lipofectamine LTX. DNA was diluted in Opti-MEM and mixed with 3.5 µL Lipofectamine LTX per well, incubated for 20 minutes, and added to cells at ∼60–70% confluency. Cells were harvested 48 hours post-transfection.

### 2-5 Oxidative Stress Assays

#### 2-5-1 Reactive Oxygen Species (ROS)

Intracellular ROS level was measured using a ROS detection kit (Canvax), based on the oxidation of 2′,7′-dichlorofluorescin diacetate (DCFH-DA), according to the manufacturer’s instructions. Cells were incubated with DCFH-DA (10µM) for 30 min at 37°C, protected from light. Following intracellular deacetylation, DCFH is oxidized by ROS to form the fluorescent compound dichlorofluorescein (DCF). Fluorescence intensity was measured by plate reader at excitation/emission wavelengths of 485/530 nm and normalized to protein content.

#### 2-5-2 Glutathione levels

Glutathione levels were measured using a Glutathione Detection Assay Kit (Cell Signaling Technology) following manufacturer instructions. Fluorescence was measured at 380/485 nm and normalized to total protein.

#### 2-5-3 Hydrogen Peroxide (H₂O₂)

H₂O₂ levels were measured using the Pierce Quantitative Peroxide Assay Kit (Thermo Fisher Scientific) according to the manufacturer’s instructions. Absorbance was measured at 560 nm, and H₂O₂ levels were normalized to total protein content.

### 2-6 Lipid Peroxidation

Lipid peroxidation was assessed using BODIPY 581/591 C11 staining (Invitrogen). Cells were incubated with 5 µM BODIPY 581/591 C11 for 60 minutes at 37°C. Reduced (non-oxidized) BODIPY 581/591 C11 was detected in the red channel, whereas oxidized BODIPY was detected in the green channel by EVOS M5000 microscope. Lipid peroxidation was expressed as the oxidized/reduced fluorescence ratio using ImageJ. Data were normalized to total cell count.

### 2-7 Fe²⁺ Measurement

Intracellular ferrous iron (Fe²⁺) level was measured using BioTracker 575 nm Fe²⁺ dye (Sigma-Aldrich). Cells were incubated with 5 µM BioTracker 575 Fe²⁺ dye for 30 minutes at 37°C and imaged by EVOS M5000 microscope. Fluorescence was quantified using ImageJ and normalized to total cell count.

### 2-8 Calcein-AM Assay

Intracellular labile iron pool (LIP) was measured using the Calcein-AM fluorescence quenching assay (Thermo Fisher Scientific). Cells were incubated with 10 µM calcein-AM for 30 minutes at 37°C followed by incubation of cells in regular medium for 1 hour. Fluorescence was quantified using ImageJ, with decreased fluorescence indicating higher intracellular iron levels due to quenching. Data were normalized to total cell count.

### 2-9 Protein Extraction Using RIPA Buffer

Cells were washed with ice-cold PBS and lysed on ice using RIPA buffer (50 mM Tris-HCl, pH 7.4, 150 mM NaCl, 1% NP-40, 0.5% sodium deoxycholate, and 0.1% SDS) supplemented with 1 mM PMSF, protease inhibitor cocktail III, and Halt phosphatase inhibitor cocktail. Cells were incubated in lysis buffer for 30 minutes on ice with occasional mixing to ensure efficient extraction. Lysates were then centrifuged at 20,000g for 15 minutes at 4°C to remove cellular debris. The resulting supernatant was collected as the total protein lysate, and protein concentration was determined using a standard protein assay. Samples were mixed with Laemmli buffer, boiled at 95°C for 5 minutes, and used for downstream Western blot analysis.

### 2-10 Western Blot Analysis

Proteins were loaded and separated by SDS–PAGE on 4-20% polyacrylamide gels (Bio-Rad) and transferred onto PVDF membranes (Bio-Rad). Membranes were blocked in Every Blot Blocking Buffer (Bio-Rad) for 30 min at room temperature and incubated overnight at 4°C with primary antibodies against targeted markers and Tubulin. Following incubation, membranes were washed and incubated with HRP-conjugated secondary antibodies for 1 hour at room temperature. Protein bands were imaged using the ChemiDoc Imaging System (Bio-Rad). Densitometric analysis was performed using Image Lab and was normalized to α-tubulin.

### 2-11 Immunofluorescence Staining

Cells were fixed with 4% paraformaldehyde for 15 minutes. Cells were permeabilized with 0.1% Triton X-100 for 10 minutes. Blocking was performed using 5% BSA for 1 hour. Cells were incubated with primary antibodies overnight at 4°C, followed by fluorescent secondary antibodies for 1 hour at room temperature. Images were acquired using EVOS M5000 microscope. Representative images were acquired at 40× magnification.

### 2-12 Antibodies

Anti-HA, rabbit monoclonal (Cell Signaling, 372), 1:1000 for WB and microscopy; anti-α-Synuclein, rabbit monoclonal (Cell Signaling Technology, #2642), 1:1000 for WB, anti-FLAG Tag, mouse monoclonal (Cell Signaling Technology, #2368), 1:1000 for WB, anti-ceruloplasmin, sheep polycolonal (BioRad, 1940-0004) 1:1000 for WB, anti-DJ rabbit monoclonal (Invitrogen), 1:1000 for WB, anti-α-synuclein, mouse monoclonal (Santa Cruz Biotechnology, clone 211, sc-12767), 1:50 for staining, anti-tyrosine hydroxylase, rabbit polyclonal (Millipore, AB152), 1:1000 for WB, anti-4-hydroxynonenal, mouse monoclonal, (R&D Systems, MAB3249) 1:1000, 76, anti-DOPA decarboxylase/DDC, mouse monoclonal (Novus Biological, clone CL2962), 1:200 for WB, anti-DAT/dopamine transporter/SLC6A3, mouse monoclonal (Santa Cruz Biotechnology, clone 6-8D6, sc-32259), 1:250 for WB, anti-mouse IgG (H+L), goat secondary antibody, Alexa Fluor™ 594 (Thermo Fisher Scientific, Highly Cross-Adsorbed), 1:200 for microscopy; anti-rabbit IgG (H+L), goat secondary antibody, Alexa Fluor™ 594 (Thermo Fisher Scientific, Highly Cross-Adsorbed), 1:200 for microscopy; anti-Transferrin Receptor/CD71, rabbit monoclonal (Cell Signaling Technology, clone D7G9X, #13113), 1:1000 for WB; anti-SLC40A1/Ferroportin-1, rabbit monoclonal (Cell Signaling Technology, clone F4A2M, #80672), 1:1000 for WB; anti-GPX4, rabbit monoclonal (Cell Signaling Technology, #52455), 1:1000 for WB; anti-FTH1, rabbit monoclonal (Cell Signaling Technology, clone D1D4, #4393), 1:1000 for WB; anti-xCT/SLC7A11, rabbit monoclonal (Cell Signaling Technology, clone D2M7A, #12691), 1:1000 for WB; anti-4F2hc/SLC3A2, rabbit monoclonal (Cell Signaling Technology, clone D3F9D, #47213), 1:1000 for WB; anti-NCOA4, rabbit monoclonal (Cell Signaling Technology, clone E8H8Z, #66849), 1:500 for WB; anti-ACSL4, rabbit monoclonal (Cell Signaling Technology, clone F6T3Z, #38493), 1:1000 for WB, anti-ceruloplasmin, rabbit polyclonal (Abcam, AB48614), 1:150 for microscopy, goat anti-rabbit, Alexa Fluor Plus 488 (ThermoFisher, A-11034); 1:600 for staining, goat anti rabbit peroxidase-conjugated secondary and goat anti mouse HRP secondary (Jackson Laboratories, 115-035-144 and 115-035-146; 1:5000) for WB.

### 2-13 Statistical Analysis

All data are presented as mean ± SD from at least three independent experiments. Statistical significance was determined using one-way or two-way ANOVA followed by Tukey’s multiple comparisons test, as appropriate. Adjusted p values < 0.05 were considered statistically significant. All analyses were conducted in GraphPad’s Prism software.

## 3 Results

### 3-1 Characterization of Stable α-Syn Expression

To establish a cellular model of α-synucleinopathy, we utilized M17 neuroblastoma cells and generated cell lines stably expressing empty vector, 1x-or 2x-α-syn. M17 cells are frequently used as a well-established dopaminergic cellular model in PD research. M17 cells retain a robust dopaminergic phenotype and are capable of robust expression of markers such as TH and DAT and highly reactive to oxidative and metabolic stress (15–17). M17 cells have also been widely used to investigate α-syn biology, dopamine-associated toxicity, and redox dysregulation relevant to neurodegenerative disease. Their compatibility with stable transgene expression and biochemical analyses also makes them a tractable system for studying the interplay between α-syn over-expression, iron remodeling, and oxidative stress pathways (18).

We utilized a pCMV-driven mammalian expression vector to establish stable cell lines. The 1x-α-syn encodes full-length human α-syn with HA-tag at the C-terminals (S1A), while 2x-α-syn was engineered by inserting a second α-syn coding sequence downstream of a self-cleaving P2A peptide (S1B). Amino acid sequence and features including cleavage site, and epitope tags are shown in Figure 1A and B. To distinguish the two products, the upstream α-syn and downstream α-syn in the 2x model had a C-terminal HA tag and a C-terminal FLAG tag, respectively (Figure 1B). Full sequences, schematic representation and features of plasmids are in S1A and S1B.

**Figure 1.**
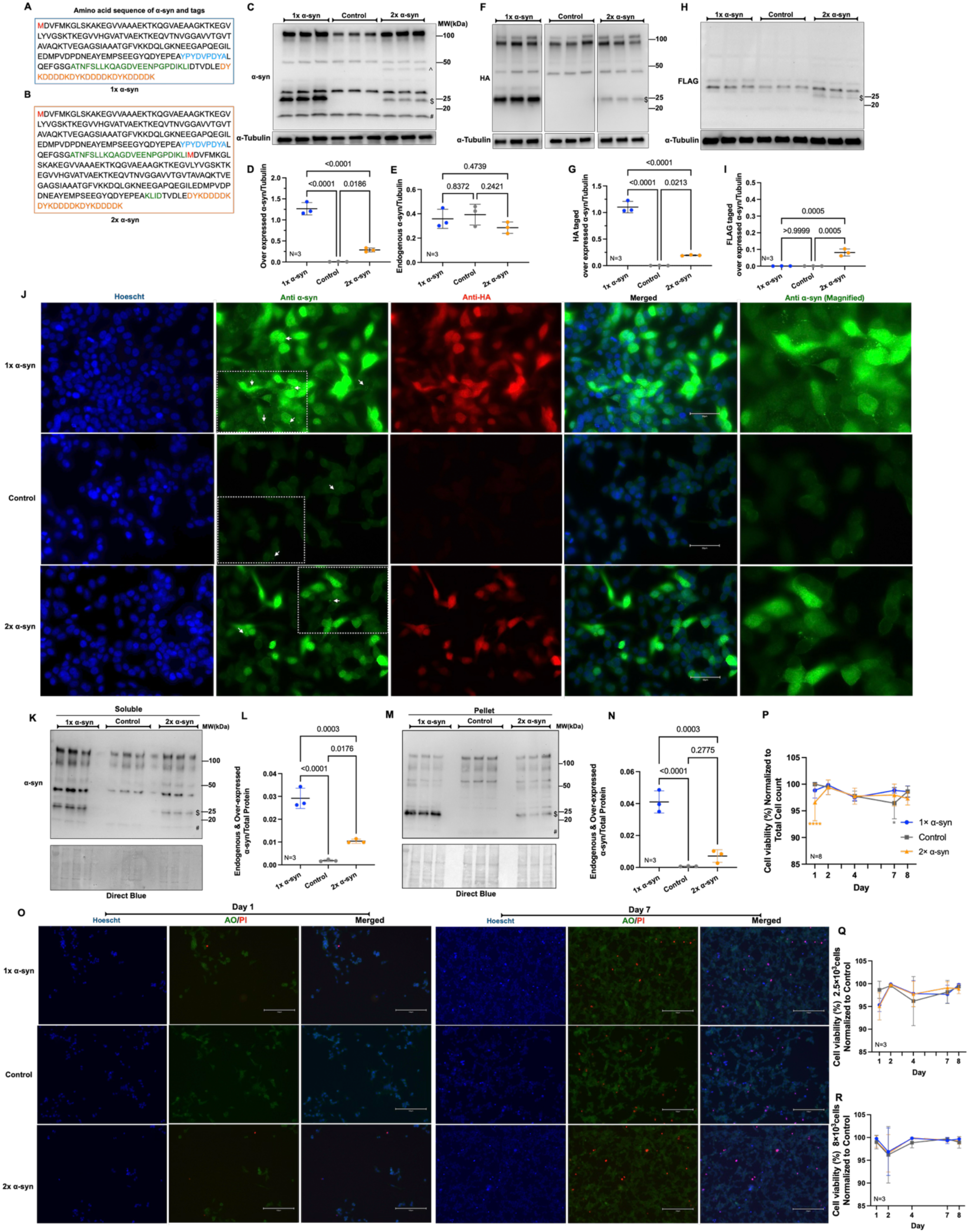
Generation and characterization of stably expressing α -synuclein M17 Cell Lines: (A-B) The amino acid sequence of the encoded human α-syn protein in 1x and 2x plasmids. Red font, starting methionine; blue font, the HA tag; green font, P2A self-cleaving sequence; orange font, FLAG tag. Full plasmid sequences are shown with annotated features in S1-A, B. (C-E) Representative Western blot and densitometric quantification of over-expressed and endogenous α-syn level normalized to α-tubulin detected by anti-α-syn antibody. $: over expressed α-syn, #: endogenous α-syn, ^: unprocessed over expressed α-syn in 2x cells. The other bands are likely non-specific bands based on panels F and G. (F-G) Western blot and densitometric quantification of over-expressed α-syn level detected by anti-HA (the original Western blot probed with HA is in S-3), (H-I) or anti-Flag. (J) Representative fluorescence images of M17 stable lines acquired at 40x magnification; green signal indicates α-syn (puncta highlighted by arrowheads), red signal indicates HA-tagged α-syn, and blue signal indicates Hoescht-stained nuclei. Scale bar = 50 μm. (K-N) Soluble and insoluble protein fractionation in stable cell lines. Representative Western blots and densitometric quantification showing α-syn distribution in (K-L) soluble and (M-N) pellet/insoluble fractions using a total α-syn antibody recognizing endogenous and exogenous protein. (O-P) AOPI viability over 8 days, showing early sensitivity in the 2x line and late-stage decline in the Control line. Representative images acquired at 20x magnification. Scale bar = 150 μm. (Q-R) Resazurin metabolic activity assays at two different seeding densities, showing stability of the adherent population. Throughout the figure, data are presented as mean ± SD from at least three independent experiments (N = 3). Statistical significance was determined using one-way or two-way ANOVA followed by Tukey’s multiple comparisons test, as appropriate. Exact p values are shown on the graphs. *p < 0.05, **p < 0.01, ***p < 0.001, ****p < 0.0001.

We validated the transgene expression in stable cell lines by Western blotting that confirmed the over-expression of α-syn (Figure 1C-I). Initially, the 2x line had higher α-syn levels than the 1x line (S2A). With continued passage, we observed reduction in α-syn expression in the 2x line (S2B). Therefore, the stable cell lines were routinely monitored, and prior to initiating the main experiments, the 1x line consistently displayed the highest overall α-syn expression. To minimize passage-dependent variation, all experiments were performed within five passages (P12-P17) and α-Syn expression was assessed at both the beginning and end of this passage range and remained stable. Based on an antibody that detects endogenous and exogenous α-syn, the 1x line expressed the highest amount of this protein compared to the Control line (which expresses the empty vector) and the 2x line (p<0.0001, p<0.0001) (Figure 1C, D). The 2x line also showed significant increase in comparison to Control (p=0.0186). When testing only for the over-expressed protein through the use of epitope tags, the 1x and 2x lines showed clear presence of α-syn with anti HA antibody, whereas the Control line showed non-specific signal (Figure 1F, G); anti-FLAG antibody only showed the over expressed α-syn in 2x line (Figure 1H-I).

Based on WBs, endogenous levels of α-syn were not impacted by over-expression of exogenous protein (Figure 1E). The fact that the 2x line exhibits lower expression levels than the 1x line with continued passaging may reflect a selective pressure against sustained high transgene expression. One possibility is that sustained expression of the 2x construct imposed greater cellular burden, resulting in selection of cells with reduced transgene expression over time (19, 20). As cells incorporated the plasmid and underwent cellular division, perhaps the 2x integration led to transgene silencing or resulted in negative selective pressure, whereas only the cells that suppressed the high-copy-number transgene survived to confluence.

Next, to visualize the localization of over-expressed α-syn in the stably expressing cells, we performed immunocytochemistry. As shown in Figure 1J, over expressed α-syn was localized in both the nucleus and cytoplasm. We observed relatively higher levels of staining for α-syn in the 1x line compared to Control and 2x; Control cells showed the lowest levels, when staining only for endogenous α-syn. We also observed instances of punctate staining in all three lines, consistent with the detection of α-syn in the pellet fraction following biochemical fractionation. For this purpose, we used soluble/pellet centrifugation protocols, in which we determined that α-syn did in fact apportion into both fractions in all three cell lines. Using an antibody recognizing total α-syn, this protein was detected in the soluble fraction across all groups (including Control cells), reflecting the presence of the endogenous protein. The 1x line exhibited the strongest soluble α-syn signal (p<0.0001 and p=0.0003 in comparison to Control and 2x, respectively) (Figure 1K, L).

In the pellet fraction of all groups, endogenous α-syn was again detectable, indicating that a portion of endogenous α-syn partitions into the insoluble fraction under basal conditions. The 1x line again showed higher levels of pelleted over-expressed α-syn, with strong signal observed using total α-syn antibody (p<0.0001 and p=0.0003 in comparison to Control and 2x, respectively). Pelleted over-expressed α-syn was also detectable in the 2x line, although to a lesser extent than in the 1x cells, and did not reach statistical significance (Figure 1M, N).

When assessing if stable over-expression of α-syn is toxic to the stable M17 cells, AOPI staining (conducted without media removal to capture cumulative cell death) in 2x line exhibited a significant initial decrease in viability on Day 1 in comparison to Control and 1x line (p<0.0001 and p=0.0133, respectively), while the Control line showed a significant (albeit mild) reduction in viability by Day 7 in comparison to 1x line (p=0.0075) (Figure 1O, P). In addition, we examined cell viability via resazurin. Following an initial recovery period post-seeding, all lines maintained high metabolic stability. Frequent media changes ensured that measurements reflected the metabolic output of the adherent, viable population, which showed no significant differences across the lines (Figure 1Q, R). Collectively, these assessments indicate that we were able to robustly detect over-expressed α-syn in M17 cells and that its stable over-expression is not consistently toxic to them.

### 3-2 Effects of α-Syn Over-expression on Ferroptosis-Associated Iron and Redox Markers

To determine whether α-syn over-expression alters cellular susceptibility to ferroptosis, we examined key regulators of this pathway, including canonical ferroptosis regulators (ACSL4, GPX4) as well as proteins controlling iron availability (FTH1) and cystine transport (4F2hc) (21–23). These factors govern ferroptotic vulnerability by regulating the balance between lipid peroxidation, antioxidant defenses, and intracellular iron availability.

Protein levels of GPX4 were comparable across all lines and ACSL4 (a critical driver of ferroptosis) showed a significant increase in 1x cells in comparison to Control and 2x (p=0.0248, p=0.0037, respectively; Figure 2B, C). In addition, α-syn expression was associated with alterations in cellular infrastructure. Ferritin heavy chain (FTH1) levels were reduced in α-syn-over-expressing cell lines, suggesting diminished iron storage capacity (p = 0.0002 and 0.0028, respectively; Figure 2D). Similarly, the System Xc subunit 4F2hc, required for stabilization of cystine transport, was decreased in 1x cells (p = 0.0126) though this change did not reach statistical significance in 2x cells (p = 0.0949; Figure 2E). These changes were consistent with reduced iron storage capacity and altered cystine transport machinery. Despite these alteration, functional analysis using BODIPY C11 fluorescence revealed that α-syn-expressing cells exhibited lower basal lipid peroxidation compared to Control (p<0.0001 and 0.0017, respectively; Figure 2F, G).

**Figure 2.**
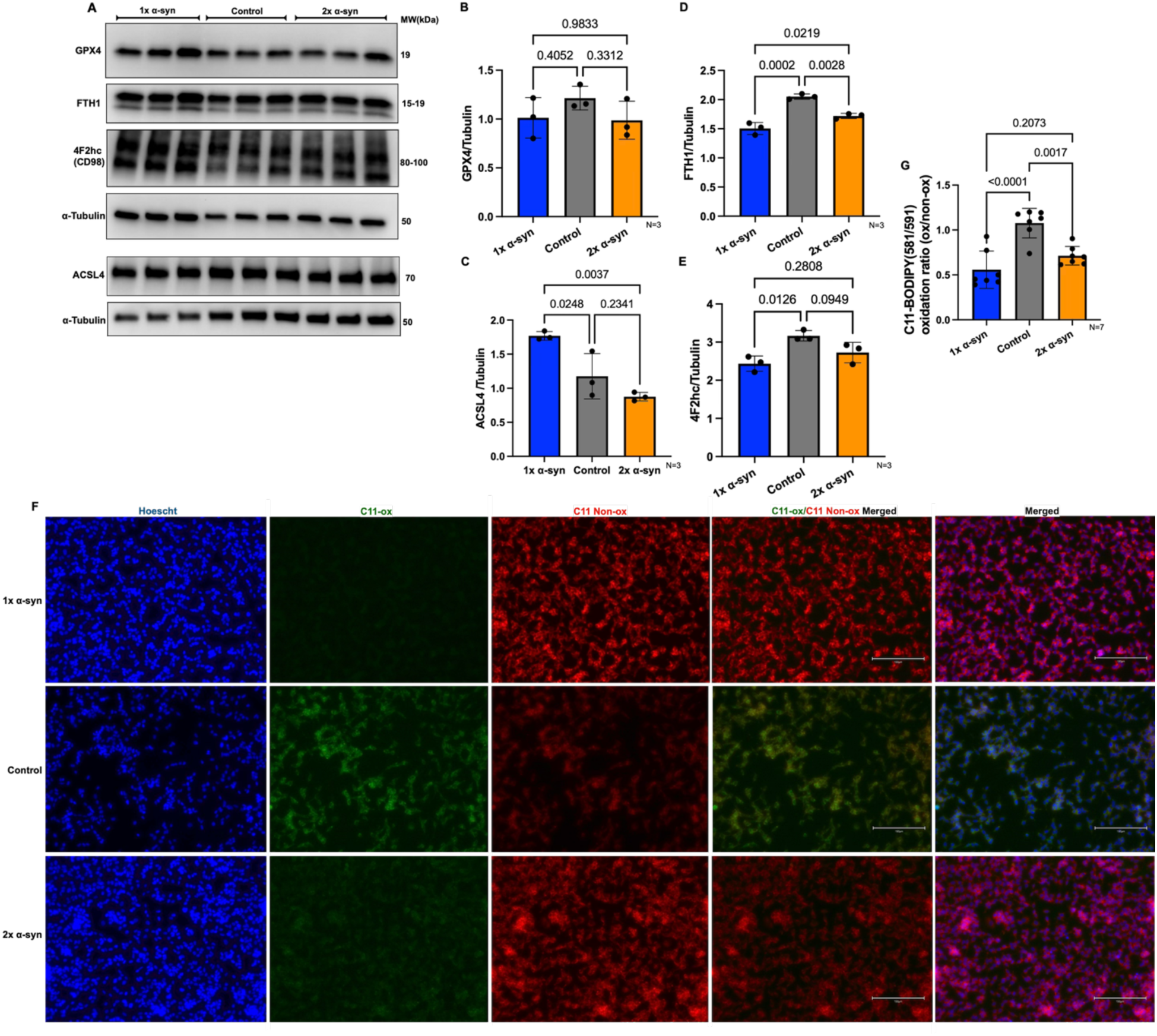
α-Syn over-expression alters iron storage and antioxidant capacity without inducing basal ferroptosis: (A) Representative immunoblots of GPX4, ACSL4, FTH1, 4F2hc, and α-Tubulin in control, 1x, and 2x α-syn-expressing M17 cell lines. (B-C) Densitometric quantification of GPX4 and ACSL4 protein levels (#: monomer ACSL4, $: dimers and multimers ACSL4). (D-E) Densitometric quantification of FTH1 and the System Xc⁻ subunit 4F2hc. (F-G) Lipid peroxidation measured by BODIPY C11 fluorescence (Normalized to total cell count). Representative images acquired at 20x magnification. Scale bar = 150 μm. Quantification was performed from at least three independent experiments. Protein levels were normalized to α-tubulin. Throughout the figure, data are presented as mean ± SD. Statistical analysis was performed using one-way ANOVA followed by Tukey’s multiple comparisons test. Exact p values are shown on the graphs. *p < 0.05, **p < 0.01, *p < 0.001, **p < 0.0001.

Based on these data, we conclude that α-syn-over-expressing cells exhibit lower basal lipid peroxidation despite significant alterations in proteins involved in iron handling and redox regulation. These observations suggest that α-syn over-expression does not induce features of a canonical ferroptotic phenotype in M17 cells. Instead, chronic α-syn expression in these cells is associated with reduced basal lipid peroxidation together with alterations in iron-handling and redox-related pathways. Therefore, α-syn-expressing cells may rely more heavily on mechanisms that regulate iron redox balance, prompting us to investigate the role of Cp.

### 3-3 Validation of Transient Ceruloplasmin Over-expression

To investigate the impact of Cp over-expression on the stable cell lines, we performed transient transfection of either Cp or of the empty pCMV vector (EV). WB confirmed robust Cp over-expression across all stable cell lines (Control, 1x, and 2x) compared to their respective EV controls (p < 0.0001). Endogenous Cp expression was nearly undetectable in the EV groups (Figure 3B, C). Cp expression levels were comparable across all three cell lines, suggesting that any subsequent differences in iron homeostasis, redox signaling, lipid peroxidation, or dopaminergic markers were not driven by unequal Cp over-expression. To determine whether Cp over-expression influenced α-syn levels, both endogenous and over-expresses α-syn were examined by Western blot. No significant changes in α-syn level were observed following Cp over-expression in any of the cell lines (Figure 3D, E). Full plasmid sequences, schematic representations, and construct features are provided in Supplementary Figure S1C.

**Figure 3.**
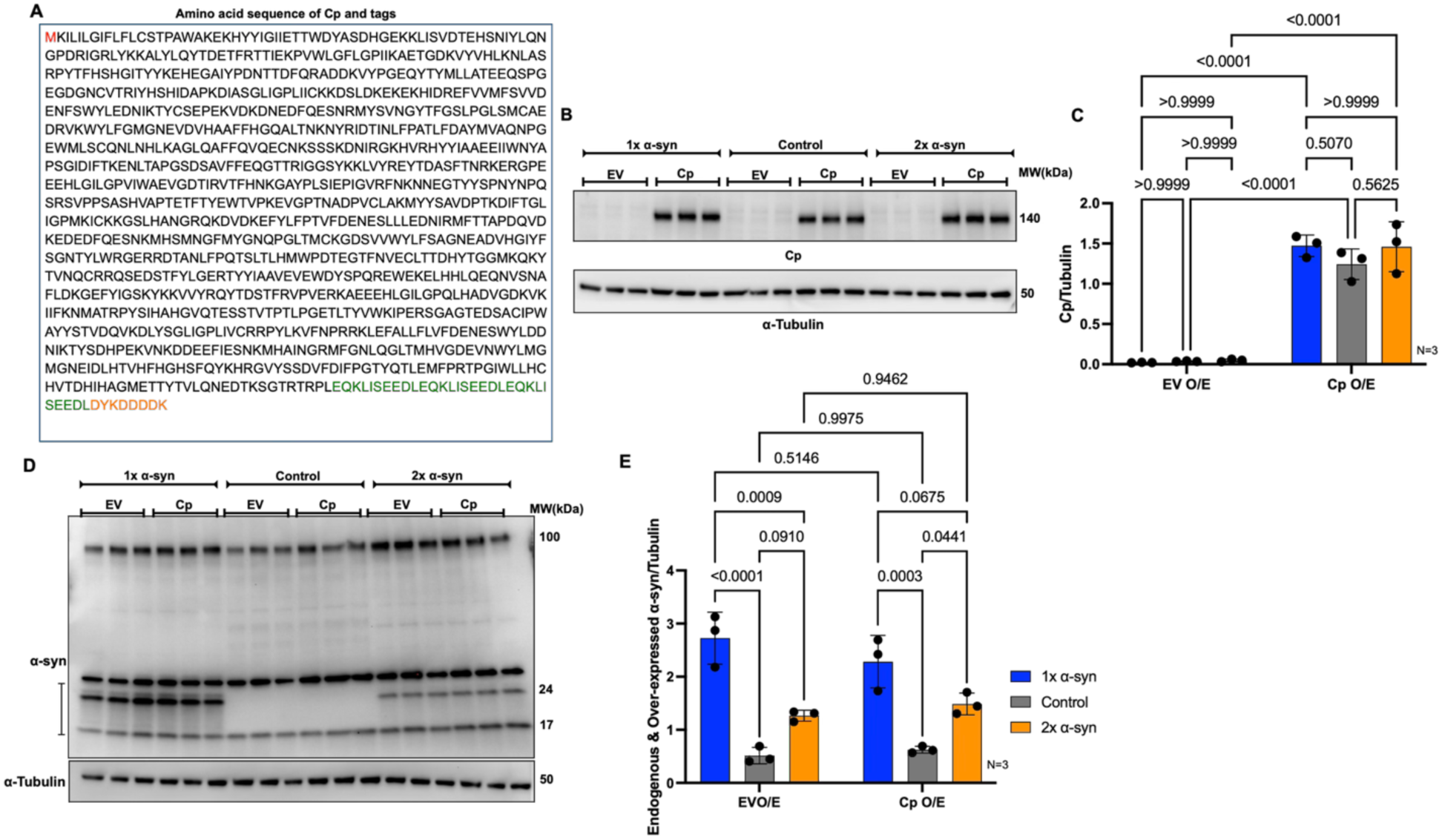
Efficient transient over-expression of Cp in M17 stable lines: (A) The amino acid sequence of the encoded human Cp protein. Red font, starting methionine; green font, 3-MYC tag; orange font, FLAG tag. Full plasmid sequences are shown with annotated features in S1-C. (B-C) Representative WB and densitometric quantification of Cp protein levels. (D-E) Western blot and densitometric quantification analysis of endogenous and exogenous α-syn levels following Cp over-expression. Protein levels were normalized to α-tubulin. Throughout the figure, data are presented as mean ± SD (n = 3 independent experiments). Statistical significance was determined using two-way ANOVA followed by Tukey’s multiple comparisons test. Exact p values are shown on the graphs. *p < 0.05, **p < 0.01, *p < 0.001, **p < 0.000.

### 3-4 Remodeling of Cellular Iron Homeostasis

To examine if α-syn and Cp over-expression collectively influence iron metabolism, we mapped the flux of intracellular iron by quantifying proteins responsible for iron import (TfR1), export (ferroportin), and storage/ferritinophagy (FTH1/NCOA4), together with functional staining for the labile iron pool (LIP) and Fe^2+^. In the EV groups, the 1x cell line maintained an iron profile that largely mirrored the Control line: no significant differences were observed in the expression of transferrin receptor 1 (TfR1), ferroportin, NCOA4, or LIP between these two groups. The only exception was FTH1, which was at significantly lower levels (p = 0.0003), which was consistent with reduced iron storage capacity (Figure 4B-F). Following Cp over-expression, the 1x remained largely the same – no significant changes were observed across most markers. The exception was a significant reduction in Fe^2+^ in 1x cells following Cp over-expression (p = 0.0031; Figure 4G), suggesting that, while the protein machinery in the 1x line was unchanged, the redox state of the existing iron pool was still subject to Cp-mediated modulation (Figure 4B-E).

**Figure 4.**
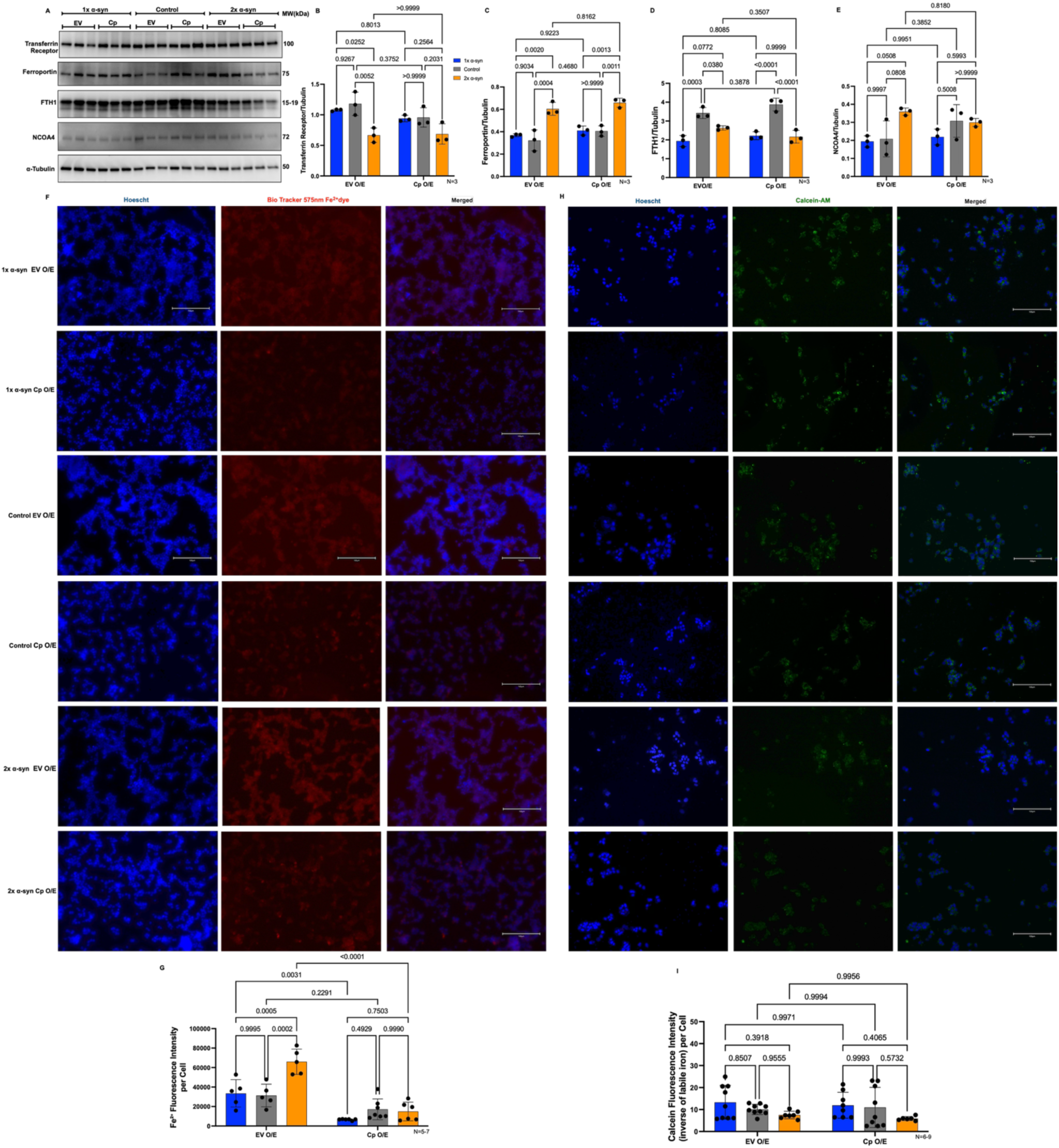
α-Syn over expression level differentially remodels iron homeostasis and response to Cp: (A) Representative WB of the level of proteins involved in iron homeostasis and α-Tubulin. (B)Transferrin Receptor 1 (TfR1), (C) Ferroportin, (D) FTH1, (E) NCOA4, (F, G) intracellular Fe²⁺ levels, and (H, I) Labile iron pool was quantified in Control, 1x, and 2x α-syn-expressing cells under EV or Cp over-expression. Throughout the figures, data are shown as mean ± SD (N=3 independent experiments, unless otherwise indicated). Statistical significance was determined by two-way ANOVA with Tukey’s multiple comparisons test (*p < 0.05, **p < 0.01, ***p < 0.001, ****p < 0.0001). Representative images acquired at 20x magnification. Scale bar = 150 μm.

In contrast to the 1x, the 2x line exhibited multiple alterations in iron-handling pathways. In EV over-expressed groups, the 2x line showed a significant decrease in TfR1 (p = 0.0052) and a significant increase in ferroportin (p = 0.0004), consistent with reduced iron import and increased iron export. (Figure 4B, C). This line also provided some evidence of active ferritinophagy: we observed lower FTH1 levels (p = 0.0380) coupled with an increase in the ferritinophagy adapter NCOA4 (p = 0.0808) (Figure 4D, 6E) (24). This mobilization of stored iron may have contributed to the significantly higher Fe^2+^ levels (p = 0.0002) found in the 2x line compared to Control (Figure 4F, G). When compared to the 1x line, the 2x (EV) line showed significant differences across all four protein markers and Fe^2+^ levels, suggesting that the 2x background exhibited distinct patterns of iron regulation (Figure 4B-G).

The effect of Cp over-expression in the 2x line was more definitive. While labile iron pool remained unchanged, Fe^2+^ levels dropped sharply (p < 0.0001), reaching levels comparable to those baseline levels observed in the Control and 1x groups (Figure 1G). This divergence where labile iron pool was comparable across comparisons but Fe^2+^ decreased provides evidence of Cp’s ferroxidase activity in converting reactive Fe^2+^ into a more stable Fe^3+^ form (25, 26). Under EV over-expression, the 1x and 2x lines showed significant differences in Fe^2+^ levels and the expression of TfR1, ferroportin, and FTH1 (p=0.0005, 0.0252, 0.0020, 0.0358, respectively; Figure 4F, G), suggesting that the two α-syn backgrounds adopt distinct patterns of iron regulation. Following Cp overexpression, most of these differences were no longer observed, although ferroportin expression remained significantly different between the two lines (p=0.0013).

Collectively, these findings indicate that consistent α-syn over-expression is associated with distinct patterns of iron regulation. The 1x line exhibited reduced FTH1 expression with relatively limited changes in other iron-related markers, whereas the 2x line showed coordinated alterations in iron import, export, and storage pathways together with elevated Fe²⁺ levels. Cp over-expression selectively reduced Fe²⁺ without significantly altering the labile iron pool, supporting a role for Cp in regulating iron redox balance rather than total intracellular iron.

### 3-5 Impact of Ceruloplasmin and α-Syn on Cellular Redox and Antioxidant Pathways

To explore whether the observed changes in iron homeostasis translated into alterations in cellular redox balance, we assessed key indicators of oxidative stress and antioxidant capacity. We first assessed intracellular ROS and H₂O₂ levels. Direct quantification of oxidative stress markers revealed that α-syn expression was not associated with increased basal ROS or H₂O₂ levels: in EV groups, both total ROS and H_2_O_2_ levels showed a non-significant decreasing trend in 1x and 2x cells compared to Control (Figure 5B, C). Cp over-expression drove a significant increase across all cell lines in comparison to their respective EV controls in ROS levels (p<0.0001; Figure 5B) and H₂O₂ level (p=0.0016 in each case; Figure 5C).

**Figure 5.**
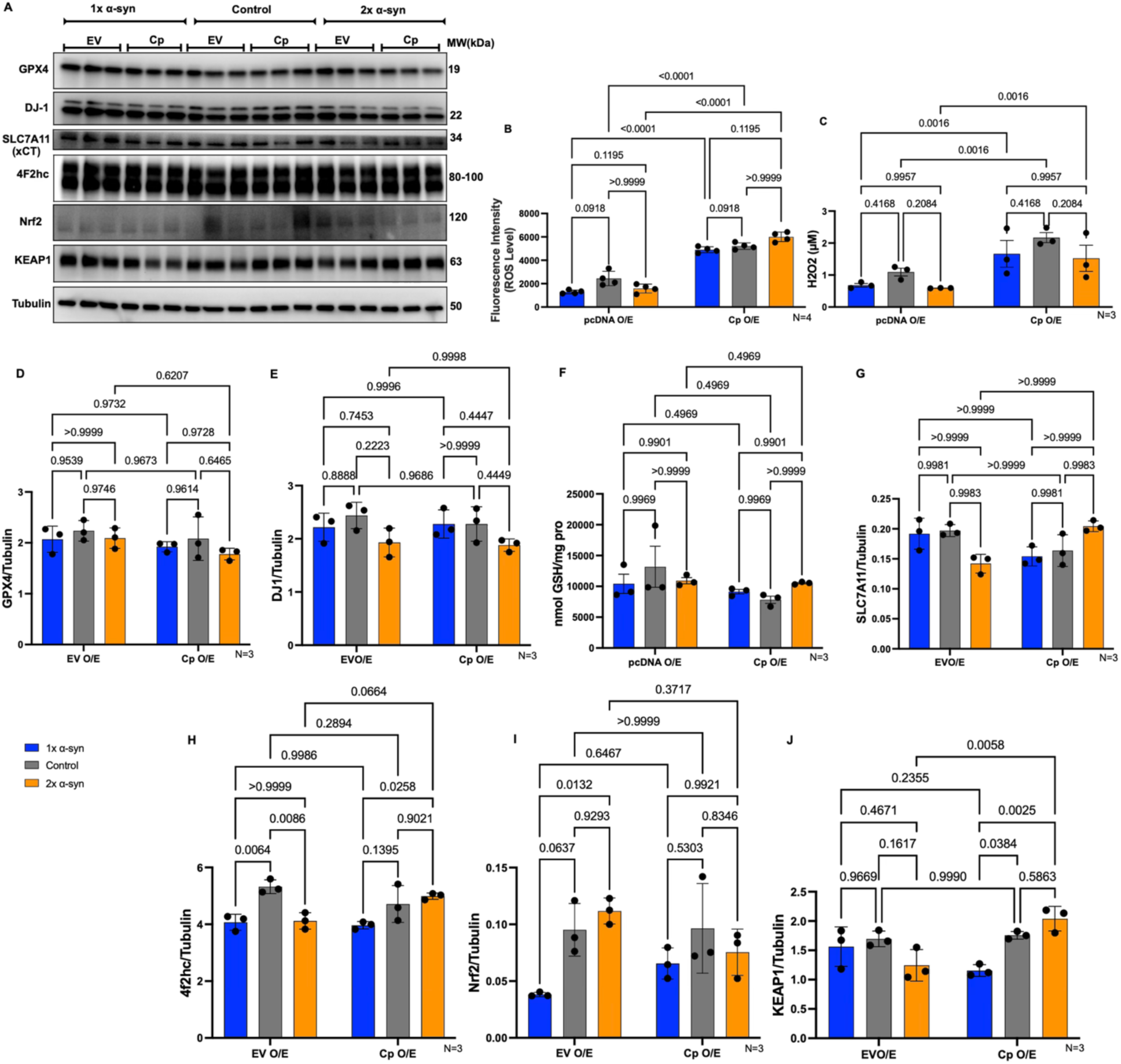
Impact of α-syn and Cp on redox transporters, stress signaling, and oxidative state: (A) Representative WB showing protein levels of GPX4, DJ-1, SLC7A11, 4F2hc, NRF2, KEAP1, and α-Tubulin in Control, 1x, and 2x α-syn stable cell lines following transient transfection with either EV or Cp expression constructs. (B–C) Intracellular oxidative stress was assessed by measuring total ROS and H₂O₂ levels. (D–E) Quantification of GPX4 and DJ-1. (F) GSH levels measurement. (G–H) Quantification of System Xc⁻ components SLC7A11 and 4F2hc. (I–J) Quantification of NRF2 and KEAP1 protein levels. Throughout the figure, data are presented as mean ± SD from three independent experiments (n = 3), unless otherwise indicated. Statistical significance was determined using two-way ANOVA followed by Tukey’s multiple comparisons test. Exact p values are shown on the graphs. *p < 0.05, **p < 0.01, *p < 0.001, **p < 0.0001.

We then investigated markers of canonical antioxidant systems: GPX4, DJ-1, and total glutathione (GSH) (27). In EV and Cp over-expression groups, GPX4 protein levels exhibited no significant difference across all groups (Figure 5D). Similarly, DJ-1 expression and total GSH pools were unaffected by either the level of the α-syn over-expression or the over-expression of Cp (Figure E, F). These data suggest that the changes in ROS and H₂O₂ were not accompanied by measurable changes in GPX4, DJ-1, or total GSH levels.

We next examined the regulation of System Xc and the Nrf2/KEAP1 signaling axis. The functional capacity of System Xc is determined by the dimerization of the catalytic subunit, SLC7A11, and the stabilizing heavy chain, 4F2hc (28). In EV controls, SLC7A11 levels were comparable among cell lines, and the over-expression of Cp did not significantly alter these levels (Figure 5G). 4F2hc protein levels showed a significant decreasing trend in EV groups, in both1x and 2x lines (p=0.0064, p=0.0086). Cp over-expression failed to restore 4F2hc in 1x line. However, a minor recovery was observed when Cp was over-expressed in the 2x line relative to its corresponding EV (Figure 5H).

We also evaluated the Nrf2/KEAP1 pathway, a central regulator of the cellular oxidative stress response (29). In EV groups, the 1x line showed a trend toward lower Nrf2 levels, whereas the 2x line showed a trend toward higher Nrf2 levels. Following Cp over-expression, Nrf2 levels were comparable across groups (Figure 5I). The negative regulator, KEAP1, remained stable in Control and 1x lines post-Cp over-expression, but increased significantly in the 2x line (0.0058). This divergent KEAP1 response shows that the 1x and 2x lines respond differently to Cp over-expression (Figure 5J). Under EV over-expression, the 1x and 2x lines did not show significant differences in ROS or H₂O₂ levels, nor in the expression of GPX4, DJ-1, GSH, 4F2hc, SLC7A11, or KEAP1. Nrf2 was the only marker different between the two lines (p = 0.0132), suggesting a possible difference in basal redox regulatory state. Following Cp over-expression, differences emerged in 4F2hc and KEAP1 expression between the 1x and 2x lines (p=0.0258, p=0.0025).

Altogether, these results suggest that chronically high α-syn levels in M17 cells are associated with lower basal ROS and H₂O₂ levels despite substantial remodeling of iron-handling pathways. These changes are not accompanied by measurable alterations in GPX4, glutathione, or DJ-1, suggesting that the reduced oxidative profile cannot be readily explained by changes in these canonical antioxidant systems. Differences in Nrf2 and KEAP1 regulation between the 1x and 2x lines, particularly following Cp over-expression, further indicate that distinct α-syn expression backgrounds exhibit different redox responses to perturbation of iron homeostasis.

### 3-6 Resistance to Lipid Peroxidation and Pro-Ferroptosis Lipid Remodeling

To investigate whether the observed remodeling of iron and redox pathways might translate into changes in lipid peroxidation, we assessed markers of lipid-derived oxidative damage and enzymatic drivers of ferroptosis. ACSL4 is a critical driver of ferroptosis and responsible for enriching cellular membranes with long-chain polyunsaturated fatty acids (PUFAs), which are highly susceptible to oxidation (21). Under EV over-expression conditions, a significant increase was observed in 1x line in comparison to Control and 2x (p=0.0432 and p=0.0037, respectively). After pro-oxidant challenge of Cp over-expression, no significant changes were detected (Figure 6A, B).

**Figure 6.**
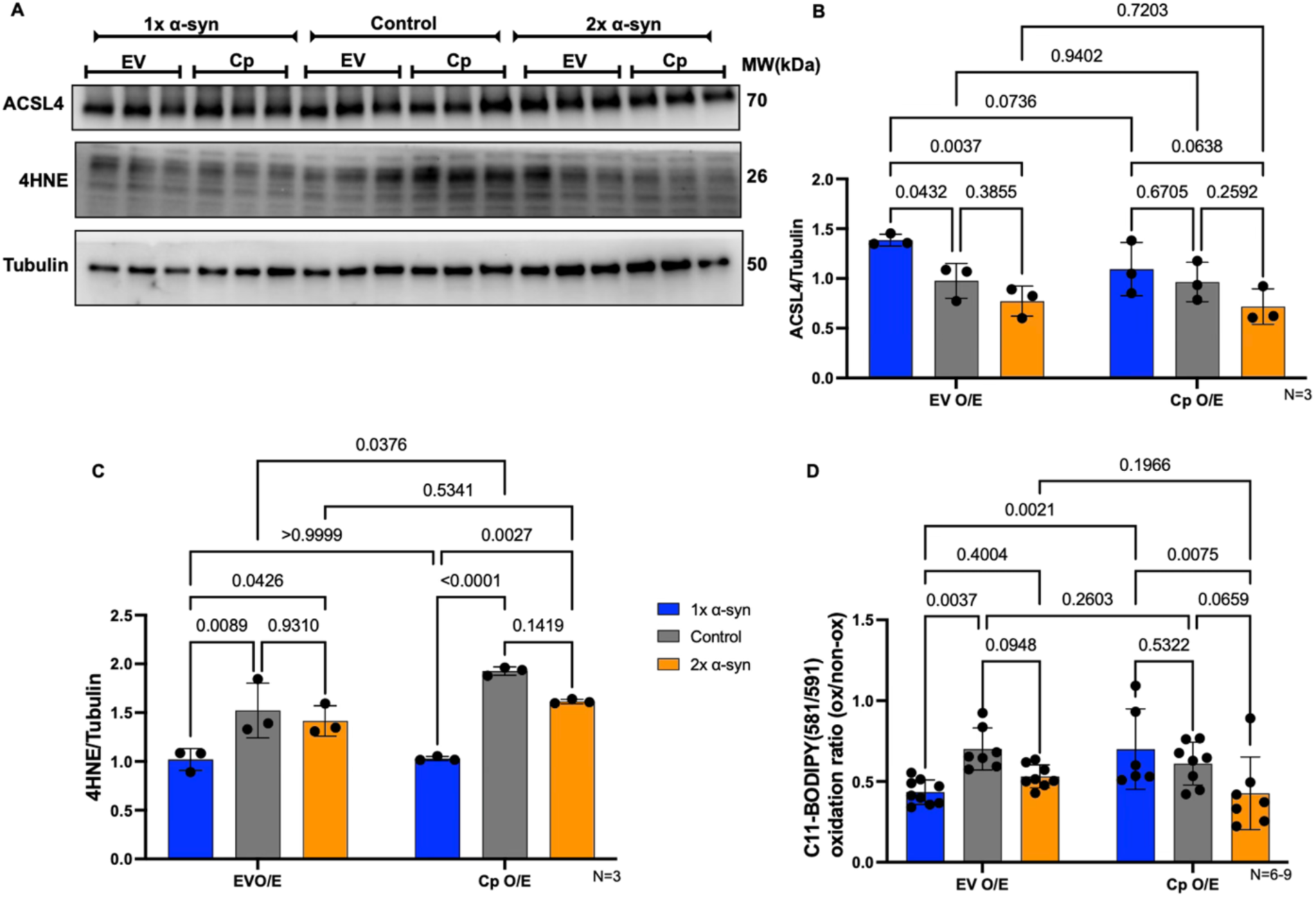
Resistance to lipid peroxidation and membrane damage: (A) Representative WB of ACSL4, 4-HNE, α-Tubulin. (B) Quantification of Western blots of the pro-ferroptotic enzyme ACSL4. (C) 4-HNE levels. (D) Quantification of BODIPY C11 Lipid Peroxidation Assay. Throughout the figure, data are presented as mean ± SD from three independent experiments (N = 3), unless otherwise indicated. Statistical significance was determined using two-way ANOVA followed by Tukey’s multiple comparisons test. Exact p values are shown on the graphs. *p < 0.05, **p < 0.01, *p < 0.001, **p < 0.0001.

We then assessed the accumulation of 4-HNE (a toxic byproduct of lipid peroxidation) and membrane-specific oxidation using the BODIPY C11 fluorescent sensor (30). In EV over-expressing groups, the 1x line exhibited significantly lower levels of 4-HNE in comparison to Control and 2x (p=0.0089, p=0.0426, respectively; Figure 6A, C). Following Cp over-expression, 4-HNE levels increased significantly in Control cells (p=0.0376), but α-syn expressing cells did not show the same response (Figure 6A, C). BODIPY C11 staining suggested that under EV over-expression, both α-syn-over-expressing lines showed lower basal lipid peroxidation compared to Control cells (p=0.0037 and p=0.0948, respectively); the greatest reduction was observed in the 1x line (Figure 6D).

Following Cp over-expression, lipid peroxidation remained largely unchanged in Control and 2x cells, while the 1x line exhibited a significant increase relative to its corresponding EV condition (p=0.0021). Under EV over-expression, the 1x and 2x lines did not show significant differences in ACSL4 expression or BODIPY C11-based oxidation readout levels, although 4-HNE levels were different between the two lines (p=0.0426). Following Cp overexpression, significant differences emerged in ACSL4 expression and lipid peroxidation measured by BODIPY C11 (p=0.0025, 0.0075), showing while differences in 4-HNE levels remained significant (p=0.0027).

These findings suggest that chronic α-syn over-expression is associated with reduced basal lipid peroxidation despite alterations in ferroptosis-associated pathways. Although ACSL4 expression is elevated in the 1x line, this is not accompanied by increased 4-HNE accumulation or BODIPY C11 oxidation under basal conditions. Furthermore, α-syn-expressing cells do not exhibit the increase in 4-HNE observed in Control cells following Cp over-expression. These data suggest that chronic α-syn expression alters the relationship between iron remodeling, oxidative stress, and lipid peroxidation, resulting in a phenotype characterized by lower basal lipid oxidative damage despite changes in several ferroptosis-related markers.

### 3-7 Dopamine Pathway Alterations Following α-Syn and Cp Over-expression

To assess any impact from α-syn and Cp over-expression on dopaminergic pathways, we examined key components of dopamine biosynthesis and markers of neuronal identity. Dopamine synthesis is primarily regulated by the rate-limiting enzyme TH and DDC (31, 32). Under EV over-expression, both the 1x and 2x α-syn lines exhibited significantly higher TH levels compared to Control cells (p = 0.0051 and p = 0.0022, respectively), while TH expression remained comparable between the 1x and 2x lines (Figure 7B). These findings suggest that α-syn expression is associated with elevated dopaminergic biosynthetic capacity. Following Cp over-expression, the 1x and 2x lines showed a further increase in TH levels relative to their EV controls (p = 0.0282 in 1x, and p = 0.0507 in 2x; Figure 7A, B). Direct comparison across Cp-over-expressed groups revealed that TH levels in the 1x and 2x lines were significantly higher than the Control (p < 0.0001; Figure 7A, B). Together, these findings suggest that sustained α-syn over-expression differentially alters dopaminergic responses to Cp-mediated oxidative challenge depending on the α-syn background.

**Figure 7.**
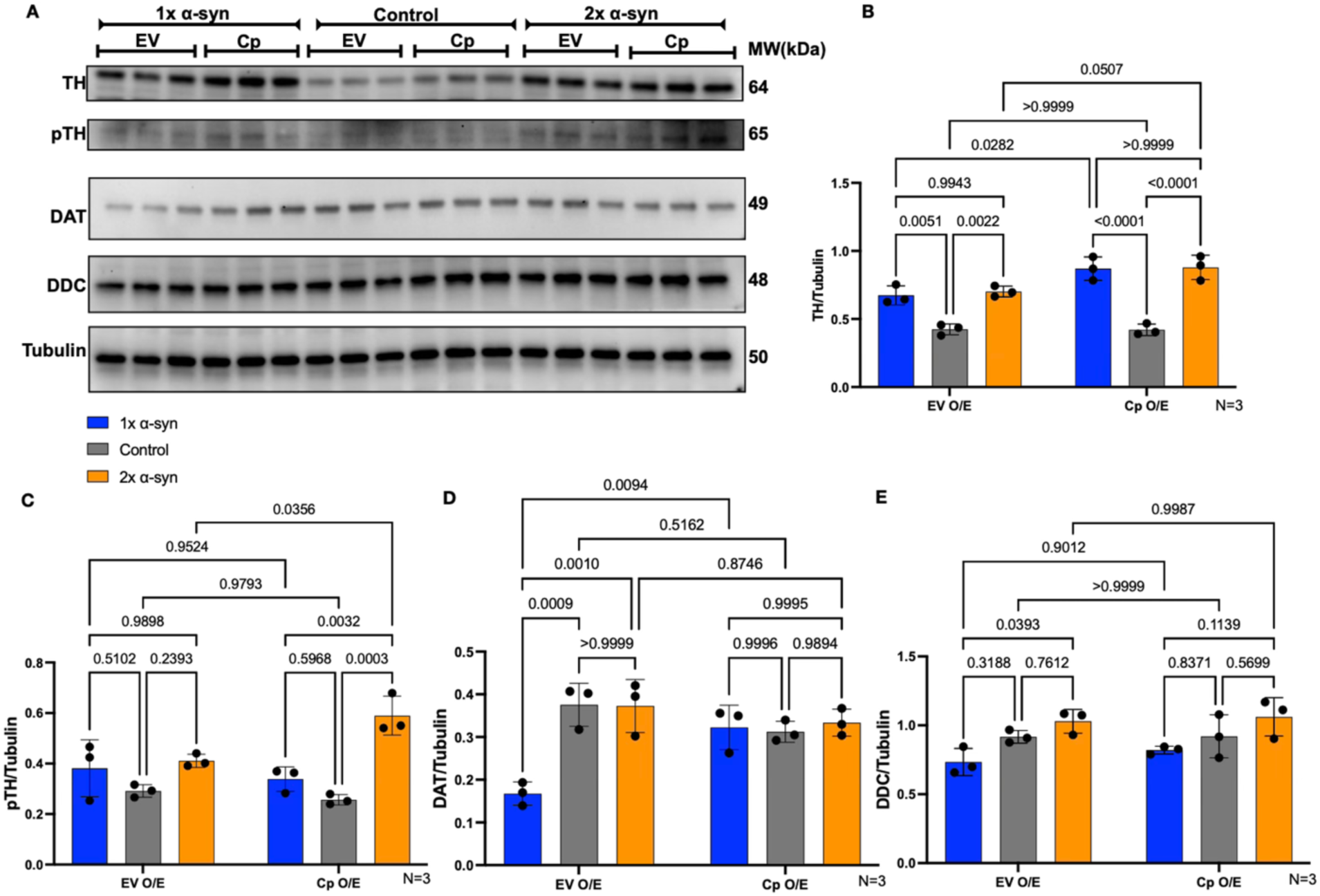
Impact on dopaminergic synthesis and identity markers: (A) Representative WB of TH, pTH, DDC, DAT and α-Tubulin in Control, 1x, and 2x α-syn stable cell lines following transient transfection with either EV or Cp expression constructs. (B) Quantification of Western blots of Total TH. (C) pTH-Ser40. (D) DDC Expression level. Bar graph showing no change under EV and Cp over-expression conditions. (E) DAT levels. Throughout the figure, data are presented as mean ± SD from three independent experiments (N = 3), unless otherwise indicated. Statistical significance was determined using two-way ANOVA followed by Tukey’s multiple comparisons test. Exact p values are shown on the graphs. *p < 0.05, **p < 0.01, *p < 0.001, **p < 0.0001.

Next, we examined phosphorylated TH at Ser40 (pTH-Ser40), an activated form of the enzyme associated with dopamine synthesis (31, 33). Under EV over-expression, pTH-Ser40 levels were comparable across all groups (Figure 7C). Following Cp over-expression, no significant changes were observed in the Control or 1x lines relative to their corresponding EV controls. In contrast, the 2x α-syn line exhibited a significant increase in pTH-Ser40 levels following Cp over-expression compared to its corresponding EV controls (p=0.0356), as well as relative to the Cp-over-expressed Control (p=0.0003) and 1x groups (p=0.0032). These findings suggest that Cp-over-expression differentially influences TH activation depending on the α-syn background.

Then, we assessed dopamine transporter (DAT) protein levels. Under EV over-expression, the 1x α-syn line exhibited significantly lower DAT levels compared to both the Control and 2x lines, while DAT expression was comparable between the Control and 2x groups (Figure 7D). Following Cp over-expression, the 1x line showed a significant increase in DAT expression relative to its corresponding EV condition. No significant changes were observed in the Control or 2x lines following Cp over-expression.

We finally examined DDC, another key enzyme involved in dopamine biosynthesis (31). Under EV over-expression, the 1x α-syn line exhibited significantly lower DDC levels compared to the 2x line (p=0.0393) while the expression was still comparable between the Control and 2x groups (Figure 7E). Following Cp over-expression, no significant differences in DDC expression were observed among the groups. Collectively, these findings suggest that α-syn over-expression is associated with distinct patterns of dopaminergic pathway regulation. While a baseline increases in TH-related signaling may reflect compensatory enhancement of dopamine biosynthetic signaling, the divergent responses observed following Cp over-expression suggest that the 1x and 2x lines differ in how dopaminergic pathways respond to perturbation of iron-redox homeostasis. The selective increase in pTH-Ser40 in the 2x line, together with the Cp-associated increase in DAT expression in the 1x line, highlights the differential regulation of dopamine synthesis and transport across the two α-syn backgrounds.

## 4 Discussion

Although PD and related synucleinopathies are characterized by the progressive accumulation of α-syn, iron dyshomeostasis, and oxidative stress, the mechanistic relationship among these processes remains incompletely understood (4, 34–37). Increasing evidence suggests that ferroptosis-associated pathways contribute to dopaminergic vulnerability, but it is unclear whether chronically higher levels of α-syn protein directly promote ferroptotic degeneration or instead alter cellular responses to iron and oxidative stress (38, 39). Here, we prepared stable M17, dopaminergic-like cell lines expressing different levels of human α-syn and used this system to probe into the interaction among α-syn expression, iron homeostasis, redox regulation, and Cp intervention. Our findings from this cellular model of high α-syn levels suggest that chronic α-syn over-expression does not induce features of a canonical ferroptotic phenotype in M17 cells. Instead, α-syn expression was associated with substantial remodeling of iron-handling pathways, reduced basal oxidative stress, lower lipid peroxidation, and distinct responses to Cp-mediated perturbation of iron redox balance.

Previous studies in postmortem PD tissue, toxin-based models, α-syn over-expressing cell models, and transgenic animals have indicated dysregulated iron metabolism often accompanied by oxidative stress and mitochondrial dysfunction (34, 37). At the same time, α-syn itself has been linked to iron metabolism through interactions with ferritin, transferrin receptor pathways, and ferroxidase-associated processes (4, 40). Reported data have further suggested that iron may accelerate α-syn aggregation and toxicity, while α-syn over-expression may sensitize neurons to oxidative injury and ferroptosis-like degeneration (36, 38). The relationship between α-syn burden, iron remodeling, and oxidative stress appears to be highly dynamic and context-dependent.

Our findings both align with and differ from previous α-syn-and ferroptosis-related models of PD. Prior works in cellular and animal systems reported that α-syn accumulation promotes oxidative stress, mitochondrial dysfunction, iron accumulation, and ferroptosis-associated degeneration (36, 38, 41). One study observed that α-syn aggregation can directly drive ferroptotic-like neuronal death through lipid peroxidation and iron-dependent oxidative injury (38). Similarly, toxin-based models, transgenic α-syn models, and viral α-syn overexpression systems have frequently associated α-syn pathology with oxidative stress and disturbances in iron homeostasis (4, 36, 42–44). Here, our over-expression model revealed a more complex phenotype in which chronic α-syn expression was associated with reduced basal ROS and lipid peroxidation despite substantial remodeling of iron-handling pathways. This discrepancy may reflect important differences between acute *versus* chronic α-syn stress, transient *versus* stable expression systems, cellular *versus* animal models of disease, and adaptive responses that emerge during long-term cellular selection. Similar adaptive phenotypes have been described in chronic neuronal stress models, where surviving populations undergo metabolic and redox reprogramming that partially suppresses oxidative burden while reducing cellular flexibility (39, 45). Our findings, therefore, suggest that chronic α-syn burden may initially induce a compensatory redox-adapted state before overt oxidative collapse occurs. In addition, our observations regarding Cp-mediated modulation of Fe^2+^ are consistent with previous studies highlighting the importance of ferroxidase activity in maintaining neuronal iron balance.

Knockout Cp Mouse models develop progressive iron accumulation and neurodegeneration, while genetic variants and oxidative modifications of Cp have been reported in PD brains and cerebrospinal fluid have been associated with impaired ferroxidase activity and disrupted iron homeostasis (46–48). These studies support the idea that disturbances in iron oxidation and export pathways contribute to neuronal vulnerability. In line with these prior conclusions, our data further suggest that Cp-mediated modulation of iron redox state may have context-dependent consequences, particularly in cells already existing within a metabolically constrained α-syn-associated state.

In contrast to the traditional view that α-syn over-expression promotes oxidative stress, our data suggest that stable α-syn expression, particularly in the 1x line, may be associated with lower basal ROS, H₂O₂, lipid peroxidation, and 4-HNE accumulation. This finding was notable because it occurred despite substantial remodeling of iron-handling pathways and alterations in proteins linked to antioxidant support. Reduced FTH1 and 4F2hc expression were consistent with changes in iron storage and cystine transport machinery, yet these alterations were not accompanied by increased basal oxidative damage. Together, these findings suggest that chronic α-syn expression can be associated with a reduced oxidative profile under basal conditions despite significant changes in iron and redox regulatory pathways (49–51).

This apparent paradox may reflect a form of long-term cellular adaptation. Chronic α-syn over-expression imposes persistent proteostatic stress on neuronal cells and may promote adaptive changes that alter how surviving populations respond to iron and oxidative stress (52, 53). The mechanisms underlying this phenotype are unclear but may involve changes in iron handling, lipid metabolism, or other stress-response pathways (54, 55). Similar adaptive responses have been reported in chronic neurodegenerative models, where prolonged cellular stress is associated with reduced oxidative burden despite ongoing pathological challenges (39, 49, 54). Our findings also differ from models in which α-syn expression directly induces a canonical ferroptotic phenotype. Canonical ferroptosis regulators, including ACSL4 and GPX4, remained largely unchanged across the stable cell lines. The relative stability of these proteins, together with reduced basal lipid peroxidation and preserved glutathione levels, suggests that chronic α-syn expression alone is insufficient to activate canonical ferroptotic pathways in M17 cells under the conditions examined here (56, 57).

At the same time, our findings do not exclude the possibility that chronic α-syn expression influences ferroptosis-associated pathways. Rather, they suggest that α-syn expression alters the relationship between iron homeostasis, oxidative stress, and lipid peroxidation without inducing features of a canonical ferroptotic phenotype. Recent studies have proposed that ferroptosis exists along a continuum and that cells may undergo partial ferroptosis-associated remodeling without overt ferroptotic cell death (39, 45). In this context, the reduced lipid peroxidation observed in our α-syn-expressing cells may reflect changes in iron-redox regulation that limit basal oxidative damage while contributing to distinct responses following secondary perturbation of iron-redox balance (38, 58).

Our findings share several features with recent study linking α-syn pathology to ferroptosis-associated pathways. Similar alterations in iron-regulatory proteins, including ferritin and ferroportin, have been reported in experimental models and human PD tissue. In addition, consistent with these reports, we did not observe substantial changes in GPX4 expression, suggesting that α-syn-associated remodeling of iron homeostasis can occur independently of major alterations in this canonical ferroptosis regulator. However, unlike study indicating increased ferroptotic vulnerability and protection by ferroptosis inhibitors, chronic α-syn expression in our model was associated with reduced basal lipid peroxidation and lower oxidative stress markers. These differences may reflect the distinct biological contexts examined by the two approaches, with ferroptosis-focused paradigms evaluating susceptibility to cell death, whereas our stable over-expression system captures cellular responses that emerge during prolonged α-syn burden (59).

One of the most notable observations in this study was the divergent behavior of the 1x and 2x α-syn lines. Although the 1x line ultimately expressed higher α-syn protein levels during downstream experiments, the 2x line exhibited more dynamic iron remodeling, including decreased TfR1, increased ferroportin, elevated NCOA4, and increased Fe^2+^ levels. This suggests that the 2x line exhibited a more pronounced iron-remodeling phenotype, potentially reflecting compensatory adaptation to long-term proteostatic stress (60–62). In contrast, the 1x line exhibited a comparatively stable iron-regulatory profile together with lower basal oxidative stress. Interestingly, the progressive reduction of α-syn expression observed in the 2x line over continued passaging likely reflects selective cellular adaptation. Stable high-level α-syn over-expression induces significant proteotoxic stress, impairs protein degradation pathways, and alter cell-cycle dynamics (53, 63). It is therefore plausible that cells maintaining excessively high transgene expression were negatively selected over time or underwent partial transgene silencing. Similar phenomena have been reported in neuronal over-expression systems where sustained expression of aggregation-prone proteins becomes incompatible with long-term proliferation and cellular stability (64, 65).

Our results also provide insight into the role of Cp within this system. Altered Cp expression and activity have been implicated in PD, aceruloplasminemia, and other neurodegenerative conditions associated with iron dysregulation (47, 66). In our model, Cp over-expression reduced Fe^2+^ levels without significantly altering labile iron pool, supporting its expected ferroxidase activity. However, Cp over-expression also acted as a potent oxidative stimulus. Despite reducing Fe^2+^, Cp induced robust increases in ROS and H_2_O_2_ across all cell lines. Remarkably, these changes were not accompanied by measurable alterations in GPX4, glutathione, or DJ-1, suggesting that the observed oxidative response occurred independently of major changes in these canonical antioxidant systems.

The α-syn-expressing lines also exhibited different responses within the Nrf2/KEAP1 and System Xc-associated pathways following Cp exposure. Although changes in Nrf2 expression were modest, the divergent trends observed between the 1x and 2x lines suggest that chronic α-syn expression may influence how antioxidant signaling pathways respond to perturbations in iron-redox balance. Nrf2 is a central regulator of antioxidant defense and cellular stress adaptation (39, 67, 68). Therefore, differences in Nrf2 and KEAP1 regulation between the two α-syn lines may contribute to the distinct oxidative responses observed following Cp treatment, where chronic α-syn expression influences not only iron homeostasis and oxidative stress but also the manner in which cells respond to secondary redox challenges (68).

α-Syn plays important roles in dopamine homeostasis, vesicular function, and regulation of dopamine-associated oxidative stress, suggesting that alterations in α-syn expression may influence neuronal vulnerability through effects on dopamine handling (69, 70). Regarding dopaminergic markers, our findings lead us to propose that chronically high α-syn levels do not simply lead to a loss of dopaminergic features but instead drives a differential remodeling of dopaminergic pathways. Under basal conditions, both α-syn-expressing lines exhibited elevated TH levels relative to Control cells, suggesting adaptive remodeling of dopamine biosynthetic pathways in response to chronic α-syn expression. Of note, the responses of the two α-syn backgrounds diverged substantially following Cp-mediated oxidative challenge. The 1x and 2x lines responded through markedly different regulatory patterns involving TH activation and DAT expression. In the 2x line, Cp exposure induced a selective increase in pTH-Ser40, suggesting enhanced activation of dopamine biosynthetic signaling pathways under oxidative stress conditions. Because TH is an iron-dependent enzyme requiring Fe²⁺ for catalytic activity, this response may be linked to the elevated labile Fe²⁺ pool and increased ferritinophagy previously observed within the 2x background (31). In this context, the combination of increased iron flux and stress-associated TH activation may reflect a a phenotype characterized by increased Fe²⁺ and enhanced pTH-Ser40 activation with heightened susceptibility to dopamine-associated oxidative stress (12, 71). In contrast, the 1x line appeared to adopt a distinct dopaminergic profile associated with lower basal oxidative stress. Despite exhibiting the highest α-syn protein burden, these cells maintained reduced basal ROS and lower labile Fe²⁺ levels while simultaneously suppressing DAT expression under baseline conditions. Because cytosolic dopamine is highly susceptible to oxidation and quinone formation, reduced DAT expression may represent an adaptive attempt to limit dopamine-associated oxidative burden during chronic α-syn stress (72, 73). Interestingly, Cp exposure partially reversed this phenotype, resulting in increased DAT expression within the 1x background. This observation suggests that despite its altered iron and redox profile, the 1x line retains a degree of capacity to respond to secondary oxidative perturbation. DDC expression remained comparatively stable across conditions, suggesting that downstream dopamine synthesis pathways are less affected than upstream regulatory components in this model. Together, these findings suggest that chronic α-syn expression is associated with distinct patterns of dopaminergic regulation that become more apparent following perturbation of iron-redox homeostasis. The selective increase in pTH-Ser40 in the 2x line and the Cp-associated increase in DAT expression in the 1x line highlight differential regulation of dopamine synthesis and transport across the two α-syn backgrounds (74–76).

We used a stable dopaminergic-like α-syn over-expression model. M17 cells retain several dopaminergic features, including TH and DAT expression, and have been used in various studies examining α-syn biology, oxidative stress, mitochondrial dysfunction, and dopamine-associated toxicity (77–80). Their compatibility with biochemical fractionation, imaging, and iron analyses makes them particularly suitable for mechanistic investigations of neurodegenerative stress pathways. Nevertheless, several limitations of the current system should be acknowledged. First, M17 cells are immortalized cells and cannot fully recapitulate the physiology of mature human dopaminergic neurons. Although these cells display dopaminergic-like characteristics, they lack the complete neuronal architecture, synaptic connectivity, and metabolic complexity present in vivo (77). Second, stable α-syn over-expression itself introduces non-physiological adaptive responses that likely differ from the gradual and heterogeneous accumulation observed in human disease (53, 63). Third, the limitation relates to the discrepancy between gene dosage and final protein expression in the 1x and 2x lines. While the 2x construct was initially designed to produce higher α-syn expression, prolonged passaging resulted in partial suppression of α-syn levels within this line. This likely reflects selective pressure or transgene silencing during long-term culture. Although this phenomenon complicated the interpretation of copy-number effects, it also provided an unexpected opportunity to compare two distinct phenotypes associated with chronic α-syn expression. Finally, while our study identified several ferroptosis-associated alterations, we did not directly induce ferroptosis using canonical ferroptotic stimuli such as erastin or RSL3, nor did we test ferroptosis inhibitors including ferrostatin-1 or liproxstatin-1. As a result, our findings are not definitive evidence that α-syn-expressing cells are fully ferroptosis-resistant. Rather, our data support the presence of ferroptosis-associated remodeling and altered susceptibility to lipid oxidative stress (39, 81, 82). Future studies should extend these findings into more physiologically relevant systems, including human iPSC-derived dopaminergic neurons, organoid models, and *in vivo* synucleinopathy models. Additional investigation into membrane lipid remodeling, mitochondrial metabolism, ferritinophagy, and NRF2 signaling dynamics may help clarify the mechanisms underlying the altered iron and redox responses observed in this study. Finally, our findings support a model in which chronic α-syn expression does not simply increase oxidative stress in a linear manner. Instead, chronic α-syn burden was associated with substantial remodeling of iron homeostasis, redox regulation, lipid peroxidation, and dopaminergic pathways. Interestingly, these changes occurred alongside reduced basal ROS, H₂O₂, and lipid peroxidation despite limited alterations in several canonical antioxidant systems. Furthermore, Cp-mediated perturbation of iron-redox balance revealed distinct responses between the 1x and 2x α-syn backgrounds, indicating that chronic α-syn expression can give rise to different patterns of cellular adaptation.

## Conclusion

These findings highlight the complex relationship between α-syn accumulation, iron regulation, oxidative stress, and dopaminergic signaling, and suggest that cellular responses to chronic α-syn burden may depend on both the extent of protein accumulation and the adaptive changes that develop over time. Together, our observations provide a framework for future studies investigating how iron oxidation, ferroptosis-associated pathways, and redox regulation contribute to dopaminergic vulnerability in PD and related synucleinopathies.

## Declarations

### Consent for publication

Not applicable

### Data Availability

Data will available upon request to the corresponding authors.

### Authorship contribution statement

M.P: Conceptualization, Methodology, Investigation, Formal analysis, Data curation, Visualization, Writing original draft. B.R: Investigation. S.P.S: Investigation. F.H.H Investigation. S.V.T: Conceptualization, Supervision, Project administration, Funding acquisition, Writing-review & editing P.A.L: Conceptualization, Supervision, Project administration, Funding acquisition, Writing-review & editing W.L.T: Conceptualization, Resources, Methodology, Writing-review & editing. All authors approved the final submitted version.

## Acknowledgments

Not applicable.

## Abbreviations

4-HNE: 4-hydroxynonenal
4F2hc: 4F2 heavy chain
ACSL4: acyl-CoA synthetase long-chain family member 4
α-syn: α-synuclein
Cp: ceruloplasmin
DAT: dopamine transporter
DDC: dopa decarboxylase
EV: empty vector
Fe²⁺: ferrous iron
Fe³⁺: ferric iron
FPN: ferroportin
FTH1: ferritin heavy chain 1
GPX4: glutathione peroxidase 4
GSH: glutathione
H₂O₂: hydrogen peroxide
KEAP1: Kelch-like ECH-associated protein 1
LIP: labile iron pool
NCOA4: nuclear receptor coactivator 4
Nrf2: nuclear factor erythroid 2-related factor 2
PD: Parkinson’s disease
pTH-Ser40: tyrosine hydroxylase phosphorylated at serine 40
PUFA: polyunsaturated fatty acid
ROS: reactive oxygen species
SLC7A11: solute carrier family 7 member 11
SNpc: substantia nigra pars compacta
System Xc⁻: cystine/glutamate antiporter system
TH: tyrosine hydroxylase
TfR1: transferrin receptor 1
WB: Western blot
DJ-1: (PARK7)
PBS: (phosphate-buffered saline)
RIPA: (radioimmunoprecipitation assay buffer)
SDS-PAGE: (sodium dodecyl sulfate–polyacrylamide gel electrophoresis)

