## Supplemental file for "Chronic α-Synuclein Over-Expression and Ceruloplasmin Challenge Promote Distinct Iron and Redox Responses in M17 Cells"

**Supplementary figure 1**

A

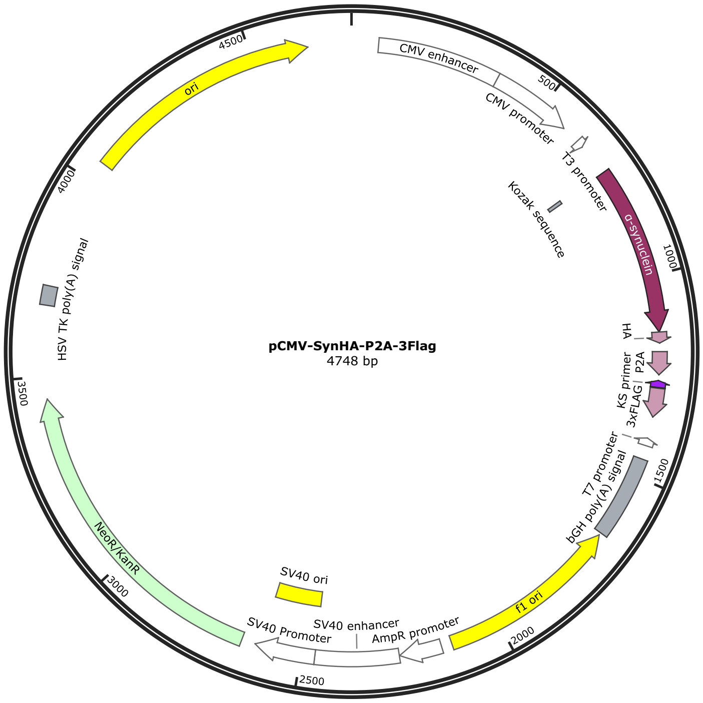


ATGCATTAGTTATTAATAGTAATCAATTACGGGGTCATTAGTTCATAGCCCATATATGGAGTT
CCGCGTTACATAACTTACGGTAAATGGCCCGCCTGGCTGACCGCCCAACGACCCCCGCCCAT
TGACGTCAATAATGACGTATGTTCCCATAGTAACGCCAATAGGGACTTTCCATTGACGTCAA
TGGGTGGAGTATTTACGGTAAACTGCCCACTTGGCAGTACATCAAGTGTATCATATGCCAAG
TACGCCCCCTATTGACGTCAATGACGGTAAATGGCCCGCCTGGCATTATGCCCAGTACATGA
CCTTATGGGACTTTCCTACTTGGCAGTACATCTACGTATTAGTCATCGCTATTACCATGGTGA
TGCGGTTTTGGCAGTACATCAATGGGCGTGGATAGCGGTTTGACTCACGGGGATTTCCAAGT
CTCCACCCCATTGACGTCAATGGGAGTTTGTTTTGGCACCAAAATCAACGGGACTTTCCAAA
ATGTCGTAACAACTCCGCCCCATTGACGCAAATGGGCGGTAGGCGTGTACGGTGGGAGGTCT
ATATAAGCAGAGCTGGTTTAGTGAACCGTCAGATCCGCTAGCGATTACGCCAAGCTCGAAAT
TAACCCTCACTAAAGGGAACAAAAGCTGGAGCTCCACCGCGGTGGCGGCCGCTCTAGCCCG
GGCGGATCCCCCGGGCTGCAGCTGCAGTCCGCCACC**ATG**GATGTATTCATGAAAGGACTTTC
AAAGGCCAAGGAGGGAGTTGTGGCTGCTGCTGAGAAAACCAAACAGGGTGTGGCAGAAGC
AGCAGGAAAGACAAAAGAGGGTGTTCTCTATGTAGGCTCCAAAACCAAGGAGGGAGTGGTG
CATGGTGTGGCAACAGTGGCTGAGAAGACCAAAGAGCAAGTGACAAATGTTGGAGGAGCA
GTGGTGACGGGTGTGACAGCAGTAGCCCAGAAGACAGTGGAGGGAGCAGGGAGCATTGCA
GCAGCCACTGGCTTTGTCAAAAAGGACCAGTTGGGCAAGAATGAAGAAGGAGCCCCACAGG
AAGGAATTCTGGAAGATATGCCTGTGGATCCTGACAATGAGGCTTATGAAATGCCTTCTGAG
GAAGGGTATCAAGACTACGAACCTGAAGCCTACCCCTACGACGTGCCCGACTACGCCCTGC
AGGAATTCGGAAGCGGAGCTACTAACTTCAGCCTGCTGAAGCAGGCTGGAGACGTGGAGGA
GAACCCTGGACCTGATATCAAGCTTATCGATACCGTCGACCTCGAGGATTACAAGGATGACG
ACGATAAGGACTATAAGGACGATGATGACAAGGACTACAAAGATGATGACGATAAATAGG
GCCCGGTACCTTAATTAATTAAGGTACCAGGTAAGTGTACCCAATTCGCCCTATAGTGAGTC

GTATTACAATTCACTCGATCGGCTCGCTGATCAGCCTCGACTGTGCCTTCTAGTTGCCAGCCA
TCTGTTGTTTGCCCCTCCCCCGTGCCTTCCTTGACCCTGGAAGGTGCCACTCCCACTGTCCTTT
CCTAATAAAATGAGGAAATTGCATCGCATTGTCTGAGTAGGTGTCATTCTATTCTGGGGGGT
GGGGTGGGGCAGGACAGCAAGGGGGAGGATTGGGAAGACAATAGCAGGCATGCTGGGGAA
CGCGTAAATTGTAAGCGTTAATATTTTGTTAAAATTCGCGTTAAATTTTTGTTAAATCAGCTC
ATTTTTTAACCAATAGGCCGAAATCGGCAAAATCCCTTATAAATCAAAAGAATAGACCGAG
ATAGGGTTGAGTGTTGTTCCAGTTTGGAACAAGAGTCCACTATTAAAGAACGTGGACTCCAA
CGTCAAAGGGCGAAAAACCGTCTATCAGGGCGATGGCCCACTACGTGAACCATCACCCTAA
TCAAGTTTTTTGGGGTCGAGGTGCCGTAAAGCACTAAATCGGAACCCTAAAGGGAGCCCCCG
ATTTAGAGCTTGACGGGGAAAGCCGGCGAACGTGGCGAGAAAGGAAGGGAAGAAAGCGAA
AGGAGCGGGCGCTAGGGCGCTGGCAAGTGTAGCGGTCACGCTGCGCGTAACCACCACACCC
GCCGCGCTTAATGCGCCGCTACAGGGCGCGTCAGGTGGCACTTTTCGGGGAAATGTGCGCGG
AACCCCTATTTGTTTATTTTTCTAAATACATTCAAATATGTATCCGCTCATGAGACAATAACC
CTGATAAATGCTTCAATAATATTGAAAAAGGAAGAATCCTGAGGCGGAAAGAACCAGCTGT
GGAATGTGTGTCAGTTAGGGTGTGGAAAGTCCCCAGGCTCCCCAGCAGGCAGAAGTATGCA
AAGCATGCATCTCAATTAGTCAGCAACCAGGTGTGGAAAGTCCCCAGGCTCCCCAGCAGGC
AGAAGTATGCAAAGCATGCATCTCAATTAGTCAGCAACCATAGTCCCGCCCCTAACTCCGCC
CATCCCGCCCCTAACTCCGCCCAGTTCCGCCCATTCTCCGCCCCATGGCTGACTAATTTTTT
TATTTATGCAGAGGCCGAGGCCGCCTCGGCCTCTGAGCTATTCCAGAAGTAGTGAGGAGGCT
TTTTTGGAGGCCTAGGCTTTTGCAAAGATCGATCAAGAGACAGGATGAGGATCGTTTCGCAT
GATTGAACAAGATGGATTGCACGCAGGTTCTCCGGCCGCTTGGGTGGAGAGGCTATTCGGCT
ATGACTGGGCACAACAGACAATCGGCTGCTCTGATGCCGCCGTGTTCCGGCTGTCAGCGCAG
GGGCGCCCGGTTCTTTTTGTCAAGACCGACCTGTCCGGTGCCCTGAATGAACTGCAAGACGA
GGCAGCGCGGCTATCGTGGCTGGCCACGACGGGCGTTCCTTGCGCAGCTGTGCTCGACGTTG
TCACTGAAGCGGGAAGGGACTGGCTGCTATTGGGCGAAGTGCCGGGGCAGGATCTCCTGTC
ATCTCACCTTGCTCCTGCCGAGAAAGTATCCATCATGGCTGATGCAATGCGGCGGCTGCATA
CGCTTGATCCGGCTACCTGCCCATTCGACCACCAAGCGAAACATCGCATCGAGCGAGCACGT
ACTCGGATGGAAGCCGGTCTTGTCGATCAGGATGATCTGGACGAAGAACATCAGGGGCTCG
CGCCAGCCGAACTGTTCGCCAGGCTCAAGGCGAGCATGCCCGACGGCGAGGATCTCGTCGT
GACCCATGGCGATGCCTGCTTGCCGAATATCATGGTGGAAAATGGCCGCTTTTCTGGATTCA
TCGACTGTGGCCGGCTGGGTGTGGCGGACCGCTATCAGGACATAGCGTTGGCTACCCGTGAT
ATTGCTGAAGAACTTGGCGGCGAATGGGCTGACCGCTTCCTCGTGCTTTACGGTATCGCCGC
TCCCGATTCGCAGCGCATCGCCTTCTATCGCCTTCTTGACGAGTTCTTCTGAGCGGGACTCTG
GGGTTCGAAATGACCGACCAAGCGACGCCCAACCTGCCATCACGAGATTTCGATTCCACCGC
CGCCTTCTATGAAAGGTTGGGCTTCGGAATCGTTTTCCGGGACGCCGGCTGGATGATCCTCC
AGCGCGGGGATCTCATGCTGGAGTTCTTCGCCCACCCTAGGGGGAGGCTAACTGAAACACG
GAAGGAGACAATACCGGAAGGAACCCGCGCTATGACGGCAATAAAAAGACAGAATAAAAC
GCACGGTGTTGGGTCGTTTGTTCATAAACGCGGGGTTCGGTCCCAGGGCTGGCACTCTGTCG
ATACCCCACCGAGACCCCATTGGGGCCAATACGCCCGCGTTTCTTCCTTTTCCCCACCCCACC
CCCCAAGTTCGGGTGAAGGCCCAGGGCTCGCAGCCAACGTCGGGGCGGCAGGCCCTGCCAT
AGCCTCAGGTTACTCATATATACTTTAGATTGATTTAAAACTTCATTTTTAATTTAAAAGGAT
CTAGGTGAAGATCCTTTTTGATAATCTCATGACCAAAATCCCTTAACGTGAGTTTTCGTTCCA
CTGAGCGTCAGACCCCGTAGAAAAGATCAAAGGATCTTCTTGAGATCCTTTTTTTCTGCGCG
TAATCTGCTGCTTGCAAACAAAAAAACCACCGCTACCAGCGGTGGTTTGTTTGCCGGATCAA

GAGCTACCAACTCTTTTTCCGAAGGTAACTGGCTTCAGCAGAGCGCAGATACCAAATACTGT
TCTTCTAGTGTAGCCGTAGTTAGGCCACCACTTCAAGAACTCTGTAGCACCGCCTACATACCT
CGCTCTGCTAATCCTGTTACCAGTGGCTGCTGCCAGTGGCGATAAGTCGTGTCTTACCGGGTT
GGACTCAAGACGATAGTTACCGGATAAGGCGCAGCGGTCGGGCTGAACGGGGGGTTCGTGC
ACACAGCCCAGCTTGGAGCGAACGACCTACACCGAACTGAGATACCTACAGCGTGAGCTAT
GAGAAAGCGCCACGCTTCCCGAAGGGAGAAAGGCGGACAGGTATCCGGTAAGCGGCAGGG
TCGGAACAGGAGAGCGCACGAGGGAGCTTCCAGGGGGAAACGCCTGGTATCTTTATAGTCC
TGTCGGGTTTCGCCACCTCTGACTTGAGCGTCGATTTTTGTGATGCTCGTCAGGGGGGCGGA
GCCTATGGAAAAACGCCAGCAACGCGGCCTTTTTACGGTTCCTGGCCTTTTGCTGGCCTTTTG
CTCACATGTTCTTTCCTGCGTTATCCCCTGATTCTGTGGATAACCGTATTACCGCC

B

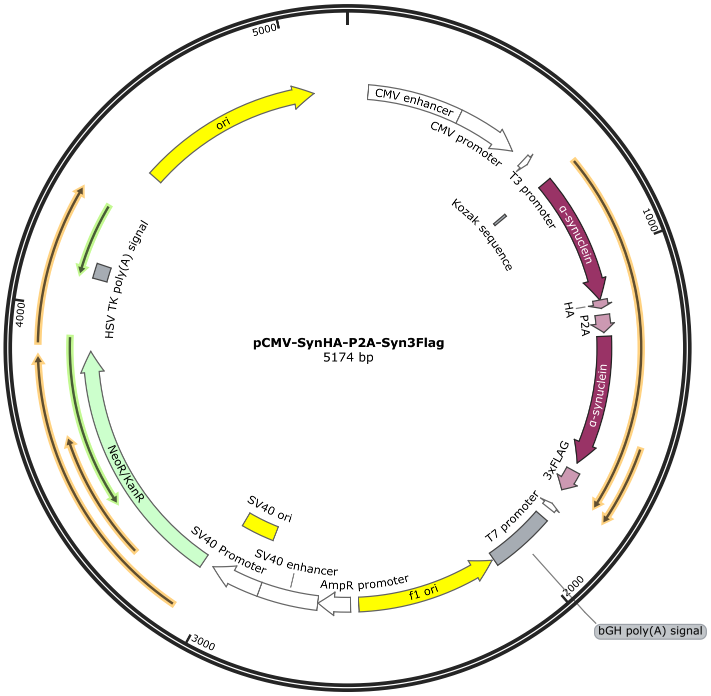


ATGCATTAGTTATTAATAGTAATCAATTACGGGGTCATTAGTTCATAGCCCATATATGGAGTT
CCGCGTTACATAACTTACGGTAAATGGCCCGCCTGGCTGACCGCCCAACGACCCCCGCCCAT
TGACGTCAATAATGACGTATGTTCCCATAGTAACGCCAATAGGGACTTTCCATTGACGTCAA
TGGGTGGAGTATTTACGGTAAACTGCCCACTTGGCAGTACATCAAGTGTATCATATGCCAAG
TACGCCCCCTATTGACGTCAATGACGGTAAATGGCCCGCCTGGCATTATGCCCAGTACATGA
CCTTATGGGACTTTCCTACTTGGCAGTACATCTACGTATTAGTCATCGCTATTACCATGGTGA
TGCGGTTTTGGCAGTACATCAATGGGCGTGGATAGCGGTTTGACTCACGGGGATTTCCAAGT
CTCCACCCCATTGACGTCAATGGGAGTTTGTTTTGGCACCAAAATCAACGGGACTTTCCAAA
ATGTCGTAACAACTCCGCCCCATTGACGCAAATGGGCGGTAGGCGTGTACGGTGGGAGGTCT
ATATAAGCAGAGCTGGTTTAGTGAACCGTCAGATCCGCTAGCGATTACGCCAAGCTCGAAAT
TAACCCTCACTAAAGGGAACAAAAGCTGGAGCTCCACCGCGGTGGCGGCCGCTCTAGCCCG
GGCGGATCCCCCGGGCTGCAGCTGCAGTCCGCCACC**ATG**GATGTATTCATGAAAGGACTTTC
AAAGGCCAAGGAGGGAGTTGTGGCTGCTGCTGAGAAAACCAAACAGGGTGTGGCAGAAGC

AGCAGGAAAGACAAAAGAGGGTGTTCTCTATGTAGGCTCCAAAACCAAGGAGGGAGTGGTG
CATGGTGTGGCAACAGTGGCTGAGAAGACCAAAGAGCAAGTGACAAATGTTGGAGGAGCA
GTGGTGACGGGTGTGACAGCAGTAGCCCAGAAGACAGTGGAGGGAGCAGGGAGCATTGCA
GCAGCCACTGGCTTTGTCAAAAAGGACCAGTTGGGCAAGAATGAAGAAGGAGCCCCACAGG
AAGGAATTCTGGAAGATATGCCTGTGGATCCTGACAATGAGGCTTATGAAATGCCTTCTGAG
GAAGGGTATCAAGACTACGAACCTGAAGCCTACCCCTACGACGTGCCCGACTACGCCCTGC
AGGAATTCGGAAGCGGAGCTACTAACTTCAGCCTGCTGAAGCAGGCTGGAGACGTGGAGGA
GAACCCTGGACCTGATATCAAGCTT**ATG**GATGTATTCATGAAAGGACTTTCAAAGGCCAAG
GAGGGAGTTGTGGCTGCTGCTGAGAAAACCAAACAGGGTGTGGCAGAAGCAGCAGGAAAG
ACAAAAGAGGGTGTTCTCTATGTAGGCTCCAAAACCAAGGAGGGAGTGGTGCATGGTGTGG
CAACAGTGGCTGAGAAGACCAAAGAGCAAGTGACAAATGTTGGAGGAGCAGTGGTGACGG
GTGTGACAGCAGTAGCCCAGAAGACAGTGGAGGGAGCAGGGAGCATTGCAGCAGCCACTG
GCTTTGTCAAAAAGGACCAGTTGGGCAAGAATGAAGAAGGAGCCCCACAGGAAGGAATTCT
GGAAGATATGCCTGTGGATCCTGACAATGAGGCTTATGAAATGCCTTCTGAGGAAGGGTATC
AAGACTACGAACCTGAAGCCAAGCTTATCGATACCGTCGACCTCGAGGATTACAAGGATGA
CGACGATAAGGACTATAAGGACGATGATGACAAGGACTACAAAGATGATGACGATAAATAG
GGCCCGGTACCTTAATTAATTAAGGTACCAGGTAAGTGTACCCAATTCGCCCTATAGTGAGT
CGTATTACAATTCACTCGATCGGCTCGCTGATCAGCCTCGACTGTGCCTTCTAGTTGCCAGCC
ATCTGTTGTTTGCCCCTCCCCCGTGCCTTCCTTGACCCTGGAAGGTGCCACTCCCACTGTCCT
TTCCTAATAAAATGAGGAAATTGCATCGCATTGTCTGAGTAGGTGTCATTCTATTCTGGGGG
GTGGGGTGGGGCAGGACAGCAAGGGGGAGGATTGGGAAGACAATAGCAGGCATGCTGGGG
AACGCGTAAATTGTAAGCGTTAATATTTTGTTAAAATTCGCGTTAAATTTTTGTTAAATCAGC
TCATTTTTTAACCAATAGGCCGAAATCGGCAAAATCCCTTATAAATCAAAAGAATAGACCGA
GATAGGGTTGAGTGTTGTTCCAGTTTGGAACAAGAGTCCACTATTAAAGAACGTGGACTCCA
ACGTCAAAGGGCGAAAAACCGTCTATCAGGGCGATGGCCCACTACGTGAACCATCACCCTA
ATCAAGTTTTTTGGGGTCGAGGTGCCGTAAAGCACTAAATCGGAACCCTAAAGGGAGCCCCC
GATTTAGAGCTTGACGGGGAAAGCCGGCGAACGTGGCGAGAAAGGAAGGGAAGAAAGCGA
AAGGAGCGGGCGCTAGGGCGCTGGCAAGTGTAGCGGTCACGCTGCGCGTAACCACCACACC
CGCCGCGCTTAATGCGCCGCTACAGGGCGCGTCAGGTGGCACTTTTCGGGGAAATGTGCGCG
GAACCCCTATTTGTTTATTTTTCTAAATACATTCAAATATGTATCCGCTCATGAGACAATAAC
CCTGATAAATGCTTCAATAATATTGAAAAAGGAAGAATCCTGAGGCGGAAAGAACCAGCTG
TGGAATGTGTGTCAGTTAGGGTGTGGAAAGTCCCCAGGCTCCCCAGCAGGCAGAAGTATGC
AAAGCATGCATCTCAATTAGTCAGCAACCAGGTGTGGAAAGTCCCCAGGCTCCCCAGCAGG
CAGAAGTATGCAAAGCATGCATCTCAATTAGTCAGCAACCATAGTCCCGCCCCTAACTCCGC
CCATCCCGCCCCTAACTCCGCCCAGTTCCGCCCATTCTCCGCCCCATGGCTGACTAATTTTTT
TTATTTATGCAGAGGCCGAGGCCGCCTCGGCCTCTGAGCTATTCCAGAAGTAGTGAGGAGGC
TTTTTTGGAGGCCTAGGCTTTTGCAAAGATCGATCAAGAGACAGGATGAGGATCGTTTCGCA
TGATTGAACAAGATGGATTGCACGCAGGTTCTCCGGCCGCTTGGGTGGAGAGGCTATTCGGC
TATGACTGGGCACAACAGACAATCGGCTGCTCTGATGCCGCCGTGTTCCGGCTGTCAGCGCA
GGGGCGCCCGGTTCTTTTTGTCAAGACCGACCTGTCCGGTGCCCTGAATGAACTGCAAGACG
AGGCAGCGCGGCTATCGTGGCTGGCCACGACGGGCGTTCCTTGCGCAGCTGTGCTCGACGTT
GTCACTGAAGCGGGAAGGGACTGGCTGCTATTGGGCGAAGTGCCGGGGCAGGATCTCCTGT
CATCTCACCTTGCTCCTGCCGAGAAAGTATCCATCATGGCTGATGCAATGCGGCGGCTGCAT
ACGCTTGATCCGGCTACCTGCCCATTCGACCACCAAGCGAAACATCGCATCGAGCGAGCACG

TACTCGGATGGAAGCCGGTCTTGTCGATCAGGATGATCTGGACGAAGAACATCAGGGGCTC
GCGCCAGCCGAACTGTTCGCCAGGCTCAAGGCGAGCATGCCCGACGGCGAGGATCTCGTCG
TGACCCATGGCGATGCCTGCTTGCCGAATATCATGGTGGAAAATGGCCGCTTTTCTGGATTC
ATCGACTGTGGCCGGCTGGGTGTGGCGGACCGCTATCAGGACATAGCGTTGGCTACCCGTGA
TATTGCTGAAGAACTTGGCGGCGAATGGGCTGACCGCTTCCTCGTGCTTTACGGTATCGCCG
CTCCCGATTCGCAGCGCATCGCCTTCTATCGCCTTCTTGACGAGTTCTTCTGAGCGGGACTCT
GGGGTTCGAAATGACCGACCAAGCGACGCCCAACCTGCCATCACGAGATTTCGATTCCACCG
CCGCCTTCTATGAAAGGTTGGGCTTCGGAATCGTTTTCCGGGACGCCGGCTGGATGATCCTC
CAGCGCGGGGATCTCATGCTGGAGTTCTTCGCCCACCCTAGGGGGAGGCTAACTGAAACAC
GGAAGGAGACAATACCGGAAGGAACCCGCGCTATGACGGCAATAAAAAGACAGAATAAAA
CGCACGGTGTTGGGTCGTTTGTTCATAAACGCGGGGTTCGGTCCCAGGGCTGGCACTCTGTC
GATACCCCACCGAGACCCCATTGGGGCCAATACGCCCGCGTTTCTTCCTTTTCCCCACCCCAC
CCCCCAAGTTCGGGTGAAGGCCCAGGGCTCGCAGCCAACGTCGGGGCGGCAGGCCCTGCCA
TAGCCTCAGGTTACTCATATATACTTTAGATTGATTTAAAACTTCATTTTTAATTTAAAAGGA
TCTAGGTGAAGATCCTTTTTGATAATCTCATGACCAAAATCCCTTAACGTGAGTTTTCGTTCC
ACTGAGCGTCAGACCCCGTAGAAAAGATCAAAGGATCTTCTTGAGATCCTTTTTTTCTGCGC
GTAATCTGCTGCTTGCAAACAAAAAAACCACCGCTACCAGCGGTGGTTTGTTTGCCGGATCA
AGAGCTACCAACTCTTTTTCCGAAGGTAACTGGCTTCAGCAGAGCGCAGATACCAAATACTG
TTCTTCTAGTGTAGCCGTAGTTAGGCCACCACTTCAAGAACTCTGTAGCACCGCCTACATACC
TCGCTCTGCTAATCCTGTTACCAGTGGCTGCTGCCAGTGGCGATAAGTCGTGTCTTACCGGGT
TGGACTCAAGACGATAGTTACCGGATAAGGCGCAGCGGTCGGGCTGAACGGGGGGTTCGTG
CACACAGCCCAGCTTGGAGCGAACGACCTACACCGAACTGAGATACCTACAGCGTGAGCTA
TGAGAAAGCGCCACGCTTCCCGAAGGGAGAAAGGCGGACAGGTATCCGGTAAGCGGCAGG
GTCGGAACAGGAGAGCGCACGAGGGAGCTTCCAGGGGGAAACGCCTGGTATCTTTATAGTC
CTGTCGGGTTTCGCCACCTCTGACTTGAGCGTCGATTTTTGTGATGCTCGTCAGGGGGGCGG
AGCCTATGGAAAAACGCCAGCAACGCGGCCTTTTTACGGTTCCTGGCCTTTTGCTGGCCTTTT
GCTCACATGTTCTTTCCTGCGTTATCCCCTGATTCTGTGGATAACCGTATTACCGCC

C

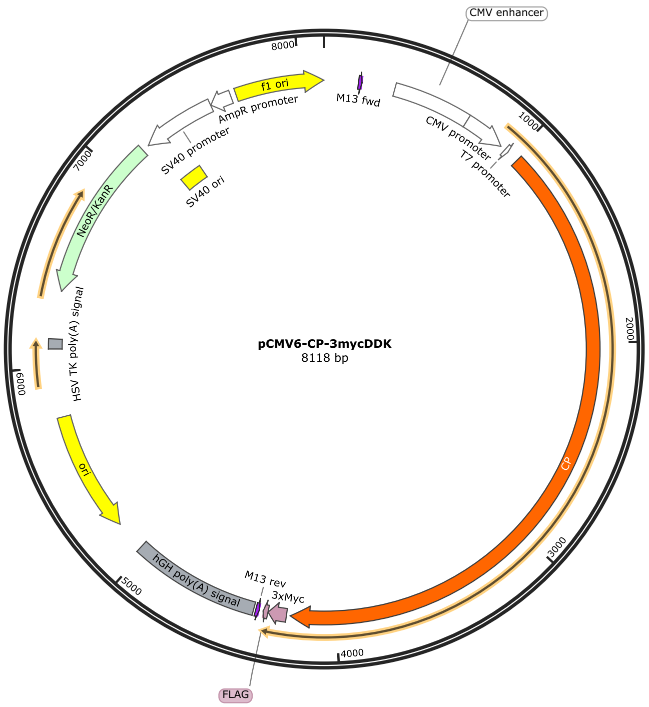


AACAAAATATTAACGCTTACAATTTCCATTCGCCATTCAGGCTGCGCAACTGTTGGGAAGGG
CGATCGGTGCGGGCCTCTTCGCTATTACGCCAGCTGGCGAAAGGGGGATGTGCTGCAAGGC
GATTAAGTTGGGTAACGCCAGGGTTTTCCCAGTCACGACGTTGTAAAACGACGGCCAGTGCC
AAGCTGATCTATACATTGAATCAATATTGGCAATTAGCCATATTAGTCATTGGTTATATAGC
ATAAATCAATATTGGCTATTGGCCATTGCATACGTTGTATCTATATCATAATATGTACATTTA
TATTGGCTCATGTCCAATATGACCGCCATGTTGACATTGATTATTGACTAGTTATTAATAGTA
ATCAATTACGGGGTCATTAGTTCATAGCCCATATATGGAGTTCCGCGTTACATAACTTACGG
TAAATGGCCCGCCTGGCTGACCGCCCAACGACCCCCGCCCATTGACGTCAATAATGACGTAT
GTTCCCATAGTAACGCCAATAGGGACTTTCCATTGACGTCAATGGGTGGAGTATTTACGGTA
AACTGCCCACTTGGCAGTACATCAAGTGTATCATATGCCAAGTCCGCCCCCTATTGACGTCA
ATGACGGTAAATGGCCCGCCTGGCATTATGCCCAGTACATGACCTTACGGGACTTTCCTACT
TGGCAGTACATCTACGTATTAGTCATCGCTATTACCATGGTGATGCGGTTTTGGCAGTACACC
AATGGGCGTGGATAGCGGTTTGACTCACGGGGATTTCCAAGTCTCCACCCCATTGACGTCAA
TGGGAGTTTGTTTTGGCACCAAAATCAACGGGACTTTCCAAAATGTCGTAATAACCCCGCCC
CGTTGACGCAAATGGGCGGTAGGCGTGTACGGTGGGAGGTCTATATAAGCAGAGCTCGTTT
AGTGAACCGTCAGAATTTTGTAATACGACTCACTATAGGGCGGCCGGGAATTCGTCGACTGG
ATCCGGTACCGAGGAGATCTGCCGCCGCGATCGCC**ATG**AAGATTTTGATACTTGGTATTTTT
CTGTTTTTATGTAGTACCCCAGCCTGGGCGAAAGAAAAGCATTATTACATTGGAATTATTGA
AACGACTTGGGATTATGCCTCTGACCATGGGGAAAAGAAACTTATTTCTGTTGACACGGAAC
ATTCCAATATCTATCTTCAAAATGGCCCAGATAGAATTGGGAGACTATATAAGAAGGCCCTT
TATCTTCAGTACACAGATGAAACCTTTAGGACAACTATAGAAAAACCGGTCTGGCTTGGGTT
TTTAGGCCCTATTATCAAAGCTGAAACTGGAGATAAAGTTTATGTACACTTAAAAAACCTTG
CCTCTAGGCCCTACACCTTTCATTCACATGGAATAACTTACTATAAGGAACATGAGGGGGCC
ATCTACCCTGATAACACCACAGATTTTCAAAGAGCAGATGACAAAGTATATCCAGGAGAGC
AGTATACATACATGTTGCTTGCCACTGAAGAACAAAGTCCTGGGGAAGGAGATGGCAATTG

TGTGACTAGGATTTACCATTCCCACATTGATGCTCCAAAAGATATTGCCTCAGGACTCATCG
GACCTTTAATAATCTGTAAAAAAGATTCTCTAGATAAAGAAAAAGAAAAACATATTGACCG
AGAATTTGTGGTGATGTTTTCTGTGGTGGATGAAAATTTCAGCTGGTACCTAGAAGACAACA
TTAAAACCTACTGCTCAGAACCAGAGAAAGTTGACAAAGACAACGAAGACTTCCAGGAGAG
TAACAGAATGTATTCTGTGAATGGATACACTTTTGGAAGTCTCCCAGGACTCTCCATGTGTG
CTGAAGACAGAGTAAAATGGTACCTTTTTGGTATGGGTAATGAAGTTGATGTGCACGCAGCT
TTCTTTCACGGGCAAGCACTGACTAACAAGAACTACCGTATTGACACAATCAACCTCTTTCC
TGCTACCCTGTTTGATGCTTATATGGTGGCCCAGAACCCTGGAGAATGGATGCTCAGCTGTC
AGAATCTAAACCATCTGAAAGCCGGTTTGCAAGCCTTTTTCCAGGTCCAGGAGTGTAACAAG
TCTTCATCAAAGGATAATATCCGTGGGAAGCATGTTAGACACTACTACATTGCCGCTGAGGA
AATCATCTGGAACTATGCTCCCTCTGGTATAGACATCTTCACTAAAGAAAACTTAACAGCAC
CTGGAAGTGACTCAGCGGTGTTTTTTGAACAAGGTACCACAAGAATTGGAGGCTCTTATAAA
AAGCTGGTTTATCGTGAGTACACAGATGCCTCCTTCACAAATCGAAAGGAGAGAGGCCCTG
AAGAAGAGCATCTTGGCATCCTGGGTCCTGTCATTTGGGCAGAGGTGGGAGACACCATCAG
AGTAACCTTCCATAACAAAGGAGCATATCCCCTCAGTATTGAGCCGATTGGGGTGAGATTCA
ATAAGAACAACGAGGGCACATACTATTCCCCAAATTACAACCCCCAGAGCAGAAGTGTGCC
TCCTTCAGCCTCCCATGTGGCACCCACAGAAACATTCACCTATGAATGGACTGTCCCCAAAG
AAGTAGGACCCACTAATGCAGATCCTGTGTGTCTAGCTAAGATGTATTATTCTGCTGTGGAT
CCCACTAAAGATATATTCACTGGGCTTATTGGGCCAATGAAAATATGCAAGAAAGGAAGTTT
ACATGCAAATGGGAGACAGAAAGATGTAGACAAGGAATTCTATTTGTTTCCTACAGTATTTG
ATGAGAATGAGAGTTTACTCCTGGAAGATAATATTAGAATGTTTACAACTGCACCTGATCAG
GTGGATAAGGAAGATGAAGACTTTCAGGAATCTAATAAAATGCACTCCATGAATGGATTCA
TGTATGGGAATCAGCCGGGTCTCACTATGTGCAAAGGAGATTCGGTCGTGTGGTACTTATTC
AGCGCCGGAAATGAGGCCGATGTACATGGAATATACTTTTCAGGAAACACATATCTGTGGA
GAGGAGAACGGAGAGACACAGCAAACCTCTTCCCTCAAACAAGTCTTACGCTCCACATGTG
GCCTGACACAGAGGGGACTTTTAATGTTGAATGCCTTACAACTGATCATTACACAGGCGGCA
TGAAGCAAAAATATACTGTGAACCAATGCAGGCGGCAGTCTGAGGATTCCACCTTCTACCTG
GGAGAGAGGACATACTATATCGCAGCAGTGGAGGTGGAATGGGATTATTCCCCACAAAGGG
AGTGGGAAAAGGAGCTGCATCATTTACAAGAGCAGAATGTTTCAAATGCATTTTTAGATAAG
GGAGAGTTTTACATAGGCTCAAAGTACAAGAAAGTTGTGTATCGGCAGTATACTGATAGCAC
ATTCCGTGTTCCAGTGGAGAGAAAAGCTGAAGAAGAACATCTGGGAATTCTAGGTCCACAA
CTTCATGCAGATGTTGGAGACAAAGTCAAAATTATCTTTAAAAACATGGCCACAAGGCCCTA
CTCAATACATGCCCATGGGGTACAAACAGAGAGTTCTACAGTTACTCCAACATTACCAGGTG
AAACTCTCACTTACGTATGGAAAATCCCAGAAAGATCTGGAGCTGGAACAGAGGATTCTGCT
TGTATTCCATGGGCTTATTATTCAACTGTGGATCAAGTTAAGGACCTCTACAGTGGATTAATT
GGCCCCCTGATTGTTTGTCGAAGACCTTACTTGAAAGTATTCAATCCCAGAAGGAAGCTGGA
ATTTGCCCTTCTGTTTCTAGTTTTTGATGAGAATGAATCTTGGTACTTAGATGACAACATCAA
AACATACTCTGATCACCCCGAGAAAGTAAACAAAGATGATGAGGAATTCATAGAAAGCAAT
AAAATGCATGCTATTAATGGAAGAATGTTTGGAAACCTACAAGGCCTCACAATGCACGTGG
GAGATGAAGTCAACTGGTATCTGATGGGAATGGGCAATGAAATAGACTTACACACTGTACA
TTTTCACGGCCATAGCTTCCAATACAAGCACAGGGGAGTTTATAGTTCTGATGTCTTTGACAT
TTTCCCTGGAACATACCAAACCCTAGAAATGTTTCCAAGAACACCTGGAATTTGGTTACTCC
ACTGCCATGTGACCGACCACATTCATGCTGGAATGGAAACCACTTACACCGTTCTACAAAAT
GAAGACACCAAATCTGGCACGCGTACGCGGCCGCTCGAGCAGAAACTCATCTCAGAAGAGG

ATCTGGAACAAAAGTTGATTTCAGAAGAAGATCTGGAACAGAAGCTCATCTCTGAGGAAGA
TCTGGATTACAAGGATGACGACGATAAGGTTTAAACGGCCGGCCGCGGTCATAGCTGTTTCC
TGAACAGATCCCGGGTGGCATCCCTGTGACCCCTCCCCAGTGCCTCTCCTGGCCCTGGAAGT
TGCCACTCCAGTGCCCACCAGCCTTGTCCTAATAAAATTAAGTTGCATCATTTTGTCTGACTA
GGTGTCCTTCTATAATATTATGGGGTGGAGGGGGGTGGTATGGAGCAAGGGGCAAGTTGGG
AAGACAACCTGTAGGGCCTGCGGGGTCTATTGGGAACCAAGCTGGAGTGCAGTGGCACAAT
CTTGGCTCACTGCAATCTCCGCCTCCTGGGTTCAAGCGATTCTCCTGCCTCAGCCTCCCGAGT
TGTTGGGATTCCAGGCATGCATGACCAGGCTCAGCTAATTTTTGTTTTTTTGGTAGAGACGGG
GTTTCACCATATTGGCCAGGCTGGTCTCCAACTCCTAATCTCAGGTGATCTACCCACCTTGGC
CTCCCAAATTGCTGGGATTACAGGCGTGAACCACTGCTCCCTTCCCTGTCCTTCTGATTTTAA
AATAACTATACCAGCAGGAGGACGTCCAGACACAGCATAGGCTACCTGGCCATGCCCAACC
GGTGGGACATTTGAGTTGCTTGCTTGGCACTGTCCTCTCATGCGTTGGGTCCACTCAGTAGAT
GCCTGTTGAATTGGGTACGCGGCCAGCGGCGAGCGGTATCAGCTCACTCAAAGGCGGTAAT
ACGGTTATCCACAGAATCAGGGGATAACGCAGGAAAGAACATGTGAGCAAAAGGCCAGCA
AAAGGCCAGGAACCGTAAAAAGGCCGCGTTGCTGGCGTTTTTCCATAGGCTCCGCCCCCCTG
ACGAGCATCACAAAAATCGACGCTCAAGTCAGAGGTGGCGAAACCCGACAGGACTATAAAG
ATACCAGGCGTTTCCCCCTGGAAGCTCCCTCGTGCGCTCTCCTGTTCCGACCCTGCCGCTTAC
CGGATACCTGTCCGCCTTTCTCCCTTCGGGAAGCGTGGCGCTTTCTCATAGCTCACGCTGTAG
GTATCTCAGTTCGGTGTAGGTCGTTCGCTCCAAGCTGGGCTGTGTGCACGAACCCCCCGTTC
AGCCCGACCGCTGCGCCTTATCCGGTAACTATCGTCTTGAGTCCAACCCGGTAAGACACGAC
TTATCGCCACTGGCAGCAGCCACTGGTAACAGGATTAGCAGAGCGAGGTATGTAGGCGGTG
CTACAGAGTTCTTGAAGTGGTGGCCTAACTACGGCTACACTAGAAGAACAGTATTTGGTATC
TGCGCTCTGCTGAAGCCAGTTACCTTCGGAAAAAGAGTTGGTAGCTCTTGATCCGGCAAACA
AACCACCGCTGGTAGCGGTGGTTTTTTTGTTTGCAAGCAGCAGATTACGCGCAGAAAAAAAG
GATCTCAAGAAGATCCTTTGATCTTTTCTACGGGGTCTGACGCTCAGTGGAACGAAAACTCA
CGTTAAGGGATTTTGGTCATGAGATTATCAAAAAGGATCTTCACCTAGATCCTTTTAAATTA
AAAATGAAGTTTTAAATCAATCTAAAGTATATATGAGTAACCTGAGGCTATGGCAGGGCCTG
CCGCCCCGACGTTGGCTGCGAGCCCTGGGCCTTCACCCGAACTTGGGGGGTGGGGTGGGGA
AAAGGAAGAAACGCGGGCGTATTGGCCCCAATGGGGTCTCGGTGGGGTATCGACAGAGTGC
CAGCCCTGGGACCGAACCCCGCGTTTATGAACAAACGACCCAACACCGTGCGTTTTATTCTG
TCTTTTTATTGCCGTCATAGCGCGGGTTCCTTCCGGTATTGTCTCCTTCCGTGTTTCAGTTAGC
CTCCCCCTAGGGTGGGCGAAGAACTCCAGCATGAGATCCCCGCGCTGGAGGATCATCCAGC
CGGCGTCCCGGAAAACGATTCCGAAGCCCAACCTTTCATAGAAGGCGGCGGTGGAATCGAA
ATCTCGTGATGGCAGGTTGGGCGTCGCTTGGTCGGTCATTTCGAACCCCAGAGTCCCGCTCA
GAAGAACTCGTCAAGAAGGCGATAGAAGGCGATGCGCTGCGAATCGGGAGCGGCGATACC
GTAAAGCACGAGGAAGCGGTCAGCCCATTCGCCGCCAAGCTCTTCAGCAATATCACGGGTA
GCCAACGCTATGTCCTGATAGCGATCCGCCACACCCAGCCGGCCACAGTCGATGAATCCAGA
AAAGCGGCCATTTTCCACCATGATATTCGGCAAGCAGGCATCGCCATGGGTCACGACGAGAT
CCTCGCCGTCGGGCATGCTCGCCTTGAGCCTGGCGAACAGTTCGGCTGGCGCGAGCCCCTGA
TGCTCTTCGTCCAGATCATCCTGATCGACAAGACCGGCTTCCATCCGAGTACGTGCTCGCTCG
ATGCGATGTTTCGCTTGGTGGTCGAATGGGCAGGTAGCCGGATCAAGCGTATGCAGCCGCCG
CATTGCATCAGCCATGATGGATACTTTCTCGGCAGGAGCAAGGTGAGATGACAGGAGATCCT
GCCCCGGCACTTCGCCCAATAGCAGCCAGTCCCTTCCCGCTTCAGTGACAACGTCGAGCACA
GCTGCGCAAGGAACGCCCGTCGTGGCCAGCCACGATAGCCGCGCTGCCTCGTCTTGCAGTTC

ATTCAGGGCACCGGACAGGTCGGTCTTGACAAAAAGAACCGGGCGCCCCTGCGCTGACAGC
CGGAACACGGCGGCATCAGAGCAGCCGATTGTCTGTTGTGCCCAGTCATAGCCGAATAGCCT
CTCCACCCAAGCGGCCGGAGAACCTGCGTGCAATCCATCTTGTTCAATCATGCGAAACGATC
CTCATCCTGTCTCTTGATCGATCTTTGCAAAAGCCTAGGCCTCCAAAAAAGCCTCCTCACTAC
TTCTGGAATAGCTCAGAGGCCGAGGCGGCCTCGGCCTCTGCATAAATAAAAAAAATTAGTC
AGCCATGGGGCGGAGAATGGGCGGAACTGGGCGGAGTTAGGGGCGGGATGGGCGGAGTTA
GGGGCGGGACTATGGTTGCTGACTAATTGAGATGCATGCTTTGCATACTTCTGCCTGCTGGG
GAGCCTGGGGACTTTCCACACCTGGTTGCTGACTAATTGAGATGCATGCTTTGCATACTTCTG
CCTGCTGGGGAGCCTGGGGACTTTCCACACCCTAACTGACACACATTCCACAGCTGGTTCTT
TCCGCCTCAGGACTCTTCCTTTTTCAATATTATTGAAGCATTTATCAGGGTTATTGTCTCATG
AGCGGATACATATTTGAATGTATTTAGAAAAATAAACAAATAGGGGTTCCGCGCACATTTCC
CCGAAAAGTGCCACCTGACGCGCCCTGTAGCGGCGCATTAAGCGCGGCGGGTGTGGTGGTT
ACGCGCAGCGTGACCGCTACACTTGCCAGCGCCCTAGCGCCCGCTCCTTTCGCTTTCTTCCCT
TCCTTTCTCGCCACGTTCGCCGGCTTTCCCCGTCAAGCTCTAAATCGGGGGCTCCCTTTAGGG
TTCCGATTTAGTGCTTTACGGCACCTCGACCCCAAAAAACTTGATTAGGGTGATGGTTCACG
TAGTGGGCCATCGCCCTGATAGACGGTTTTTCGCCCTTTGACGTTGGAGTCCACGTTCTTTAA
TAGTGGACTCTTGTTCCAAACTGGAACAACACTCAACCCTATCTCGGTCTATTCTTTTGATTT
ATAAGGGATTTTGCCGATTTCGGCCTATTGGTTAAAAAATGAGCTGATTTAACAAAAATTTA
ACGCGAATTTT

Supplementary figure 1: (A, B) Full sequences, schematic representation and features of plasmids encoding one copy and two copy of human α-syn protein. Red bold font, starting methionine; blue font, the HA tag; green font, P2A self-cleaving sequence; orange font, FLAG tag. (C) Full sequence, schematic representation and features of plasmids encoding human Ceruloplasmin. Red bold font, starting methionine; green font, 3-MYC tag; orange font, FLAG tag.

**Supplementary figure 2**

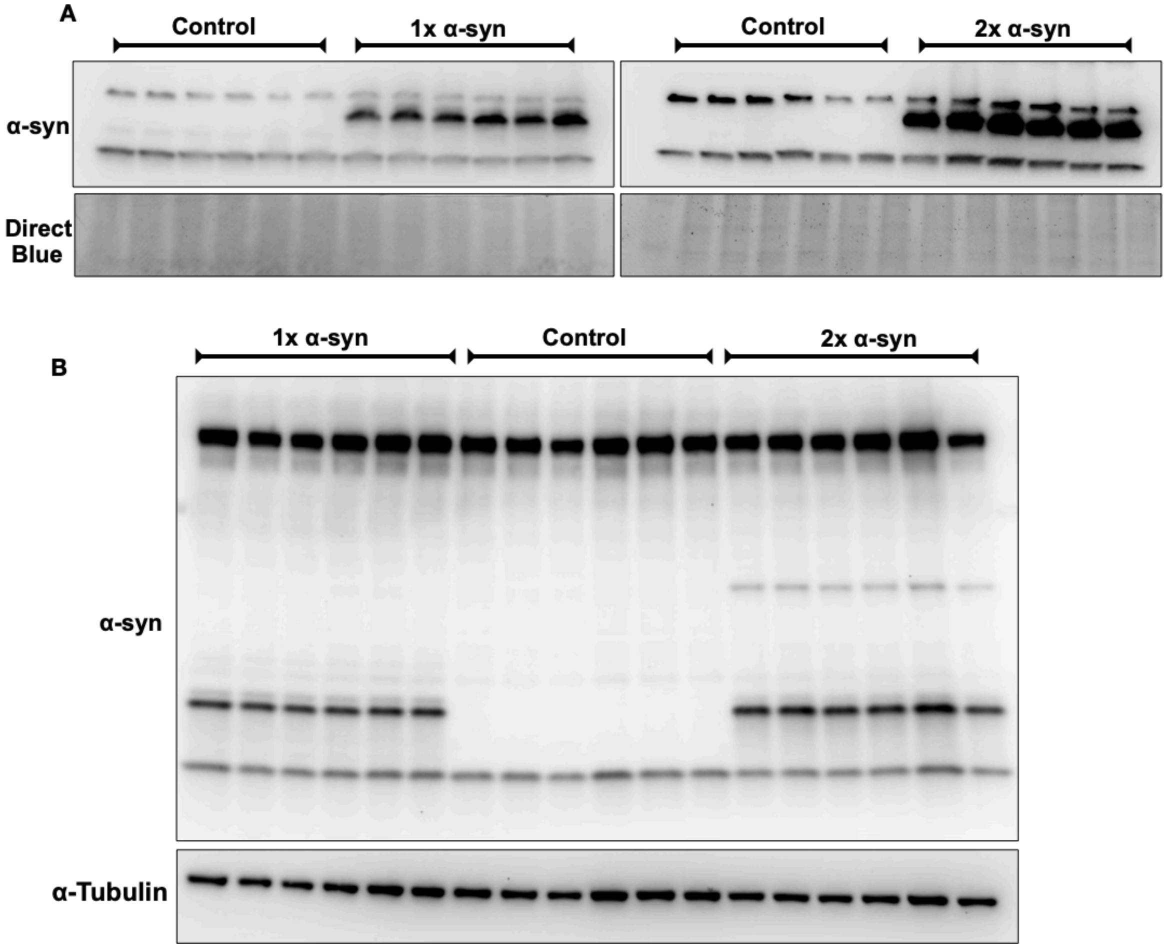


Supplementary 2: (A) Representative Western blot of over-expressed and endogenous α-syn in stable cell lines comparing α-synuclein levels in Control, 1x, and 2x cell lines at the start of the stable cell lines establishment and (B) after 3 passages (P5-P8). The blot highlights how α-synuclein expression changes over time in 2x line. Direct Blue and tubulin to compare protein loading in lanes.

**Supplementary figure 3**

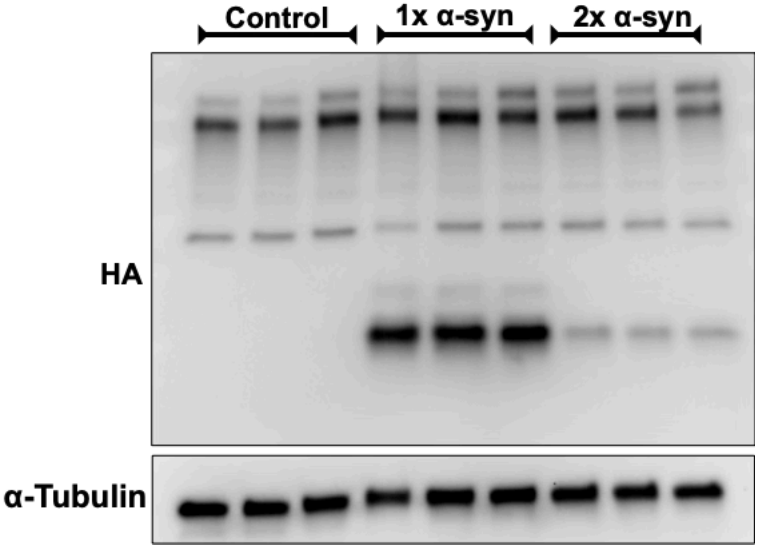


Supplementary figure 3: The original blot of over-expressed α-syn level in cell lines detected by anti-HA (Figure 1F).
